# Sterols lower energetic barriers of membrane bending and fission necessary for efficient clathrin mediated endocytosis

**DOI:** 10.1101/2021.01.31.428633

**Authors:** Ruthellen H. Anderson, Kem A. Sochacki, Harika Vuppula, Brandon L. Scott, Elizabeth M. Bailey, Maycie M. Schultz, Jason G. Kerkvliet, Justin W. Taraska, Adam D. Hoppe, Kevin R. Francis

## Abstract

As the principal internalization mechanism in mammalian cells, clathrin-mediated endocytosis (CME) is critical for cellular signal transduction, receptor recycling, and membrane homeostasis. Acute depletion of cholesterol disrupts CME, motivating analysis of CME dynamics in the context of disrupted cholesterol synthesis, sterol specificity, mechanisms involved, and relevance to disease pathology. Using genome-edited cell lines, we demonstrate that inhibition of post-squalene cholesterol biosynthesis as observed in inborn errors of cholesterol metabolism, results in striking immobilization of CME and impaired transferrin uptake. Imaging of membrane bending dynamics and CME pit ultrastructure revealed prolonged clathrin pit lifetimes and accumulation of shallow clathrin-coated structures that scaled with diminishing sterol abundance. Moreover, fibroblasts derived from Smith-Lemli-Opitz syndrome subjects displayed reduced CME function. We conclude that sterols lower the energetic costs of membrane bending during pit formation and vesicular scission during CME and suggest reduced CME contributes to cellular phenotypes observed within disorders of cholesterol metabolism.

## INTRODUCTION

Lipid composition is critical to the biophysical properties of membrane bilayers, permitting maintenance of essential cellular functions such as membrane trafficking and signal transduction (Kusumi et al., 2012; Raghunathan and Kenworthy, 2018). Cholesterol has long been recognized as a major regulator of lipid organization. Its unique structure influences lipid packing, protein localization and interactions, and global membrane properties including permeability, fluidity, and rigidity (Maxfield and Tabas, 2005). Synthesized *de novo* in the endoplasmic reticulum or derived from circulating low-density lipoproteins, unesterified cholesterol is distributed heterogeneously amongst various cell membranes, being predominately enriched at the plasma membrane (PM) where it accounts for up to 45% of the total lipids present (Das et al., 2013) and 60-90% of total cellular cholesterol (de Duve 1971; Lange et al., 1989). The importance of cholesterol composition to PM structure and function is reflected in the tight regulation of intracellular transport and highly coordinated synthetic and efflux mechanisms maintaining physiologic cholesterol levels (Luo et al., 2019; Mitsche et al., 2015; Nohturfft et al., 1998; Sun et al., 2005; Tontonoz and Mangelsdorf, 2003; Vedhachalam et al., 2007).

Membrane properties including rigidity and tension are especially pertinent to highly dynamic membrane remodeling processes, such as endocytosis (Hassinger et al., 2017). As the major pathway regulating receptor-mediated signaling, clathrin-mediated endocytosis (CME) mediates nutrient uptake, signal transduction, receptor recycling and desensitization, membrane homeostasis, and developmental regulation (Kaksonen and Roux, 2018). Evidence for cholesterol’s role in CME has been demonstrated by acute stripping of membrane cholesterol using methyl-beta-cyclodextrin (MβCD) (Rodal et al., 1999; Subtil et al., 1999). Although MβCD is a relatively non-specific means of cholesterol extraction contingent upon the carrier’s hydrophobic cavity (Zidovetzki and Levitan, 2007), these classic experiments suggest a strong defect in clathrin-coated pit budding and internalization upon MβCD treatment.

Amidst the extensive CME machinery described to date, the influence of cholesterol on plasma membrane architecture may be overlooked as an active component within assembly, curvature generation, and scission of clathrin-coated pits. CME is initiated by the recruitment of clathrin triskelia by adapter and accessory proteins to sites of phosphatidylinositol-4,5-bisphosphate (PtdIns(4,5)P_2_) enriched PM (Cocucci et al., 2012). This assembly process may be sensitive to alterations of lipid packing or shielding of charges secondary to cholesterol availability (Hirama et al., 2017). Progression of the PM from flat to highly curved clathrin-coated vesicles also requires overcoming stiffness of the membrane and tension generated by lateral membrane recruitment and opposed by membrane-cytoskeletal adhesions (Bucher et al., 2018; Scott et al., 2018). Recruitment of epsin NH2-terminal homology (ENTH), AP180 N-terminal homology (ANTH) and later Bin/Amphiphysin/Rvs (BAR) domain containing proteins likely contribute to the energy required for membrane bending (Liu et al., 2009; McMahon and Gallop, 2005; Miller et al., 2015). By interacting with these curvature effectors and stabilizing positive PM fluctuations, the clathrin lattice is thought to act as a Brownian ratchet promoting curvature formation (Liu et al., 2009). This remodeling process creates energetically unfavorable membrane stress which may be alleviated by cholesterol through rapid transbilayer redistribution (Bruckner et al., 2009). During vesicular scission, an energetically costly constriction requiring steep membrane curvature must be achieved. The GTPase dynamin catalyzes this event by polymerizing into a helical coat and constricting the endocytic pit neck by GTP hydrolysis and removal of dynamin subunits. However, previous work has demonstrated constriction alone is insufficient for scission to occur (Danino et al., 2004; Roux et al., 2006). Considering the severity of membrane bending at these sites, the mechanical properties of the PM immediately adjacent to the dynamin coat, including tension and bending rigidity, are thought to influence scission (Morlot et al., 2012). Bursts of actin polymerization also aid in overcoming resistance to membrane bending by providing additional force leading to vesicle internalization and scission (Collins et al., 2011; Grassart et al., 2014), especially in areas of high membrane tension (Boulant et al., 2011).

Human diseases associated with disruption of cholesterol homeostasis may provide insight into the lipid-specific requirements of CME, and conversely, find application from an understanding of cellular trafficking within these disease states. The best characterized of these orphan diseases, Smith-Lemli-Opitz syndrome (SLOS), is caused by genetic mutations within the post-squalene cholesterol synthesis enzyme 3β-hydroxy-steroid-Δ^7^-reductase (DHCR7) (Wassif et al., 1998) and presents clinically with multiple congenital malformations including distinctive facial features, cardiac defects, genitourinary abnormalities, syndactyly, gastrointestinal intolerance, and profound CNS dysfunction (Smith et al., 1964). Clinical severity generally correlates inversely with residual enzymatic activity; however, significant phenotypic variability exists even amongst similar pathogenic variants (Cunniff et al., 1997; Wassif et al., 2005; Waterham and Hennekam, 2012), which ranges on a wide continuum from mild cognitive impairment to embryonic lethality (Tint et al., 1995). The relative contribution of cholesterol depletion and/or accumulation of sterol precursors such as 7-dehydrocholesterol (7DHC) to disease pathogenesis also remains an open question (Wassif et al., 2017). To date, membrane defects within SLOS have primarily been attributed to disruption of lipid rafts (Gou-Fabregas et al., 2016; Keller et al., 2004). As a theory to explain membrane heterogeneity, the propensity for cholesterol to tightly pack sphingomyelin and less saturated phospholipids underlies the basis for liquid-ordered (*L*_o_) and liquid-disordered (*L*_d_) phase separation. As such, cholesterol depletion is often considered evidence for lipid raft involvement. Nevertheless, consideration of global effects on plasma membrane architecture is warranted, which may perturb processes that are themselves independent of lipid rafts (Kwik et al., 2003; Sengupta et al., 2007; van Rheenen et al., 2005). While sterols may influence developmental pathways through a variety of mechanisms, one hypothesis suggests that SLOS pathogenesis manifests in a highly temporal and tissue specific context, driven by changes in signal transduction pathways (Tulenko et al., 2006). Thus, defining the effects of disrupted sterol synthesis on the major pathway regulating receptor-mediated signaling may also provide insight into the molecular and cellular basis for the physiologic manifestations within these rare diseases.

In this study, we demonstrate that multiple stages of membrane remodeling during CME progression are sensitive to sterol-dependent membrane properties. Clathrin pit ultrastructure with cholesterol depletion exhibited asymmetric distorted structures, consistent with aberrant membrane bending trajectories, suggesting that clathrin lattice assembly competes with the elevated free energy costs of bending. Using sterol substitution strategies, we further demonstrate that a small, C3 polar headgroup and B-ring conformation constitute structural requirements for supporting efficient CME. Finally, we present evidence for disruption of CME dynamics within SLOS and conclude that sterol abundance lowers the energetic barrier for curvature generation during initiation of membrane bending and formation of the vesicle neck during clathrin-mediated endocytosis.

## RESULTS

### Clathrin-mediated endocytosis is inhibited by altered cholesterol homeostasis

To test the hypothesis that sterol homeostasis contributes to efficient CME, we evaluated CME during acute cholesterol depletion and perturbations of cholesterol synthesis in HEK293T cells. Endogenous cholesterol biosynthesis was induced by culturing cells in media containing 7.5% lipoprotein-deficient fetal bovine serum (LPDS). We used multiple targeting strategies to delineate the effects of cholesterol depletion versus terminal cholesterol precursor accumulation within the competing Bloch and Kandutsch-Russell (KR) pathways (**Figure 1A**). AY9944 dihydrochloride, a potent inhibitor of DHCR7 (Δ7-sterol reductase) (Fernandez et al., 2005; Gaoua et al., 1999; Rahier and Taton, 1996), has been used extensively to recapitulate the biochemical hallmarks of SLOS both *in vitro* and *in vivo* (Francis et al., 2016; Kolf-Clauw et al., 1996). U18666A, a potent noncompetitive inhibitor of DHCR24, was used to inhibit the final step in the Bloch pathway at nanomolar concentrations where it does not perturb cholesterol trafficking (Cenedella, 2009; Lu et al., 2015). Comparison of AY9944 (KR targeting) with U18666A (Bloch targeting) also provided access to structure-activity relationships. Specifically, DHCR7 mediated reduction of the C(7-8) double bond confers the linear cholesterol ring structure (Serfis et al., 2001), such that treatment with AY9944 leads to accumulation of non-planar sterols. Conversely, inhibition of DHCR24 leads to accumulation of sterols with more rigidity as it prevents the DHCR24 mediated reduction of C(24-25) (Chen and Tripp, 2012). The HMG-CoA reductase inhibitor Atorvastatin was used to inhibit synthesis of cholesterol and other biosynthetic intermediates, while MβCD was used to acutely deplete sterols from the plasma membrane (Rodal et al., 1999; Subtil et al., 1999). In this study, sterol content in HEK293T cells relative to LPDS controls was raised ~25% when cultured in FBS and reduced by ~50% with Atorvastatin, AY9944, or MβCD. AY9944 treatment effectively led to the predicted accumulation of 7-dehydrocholesterol and zymostenol, while U18666A induced a mild accumulation of desmosterol (**Figure 1B**).

**Figure 1.**
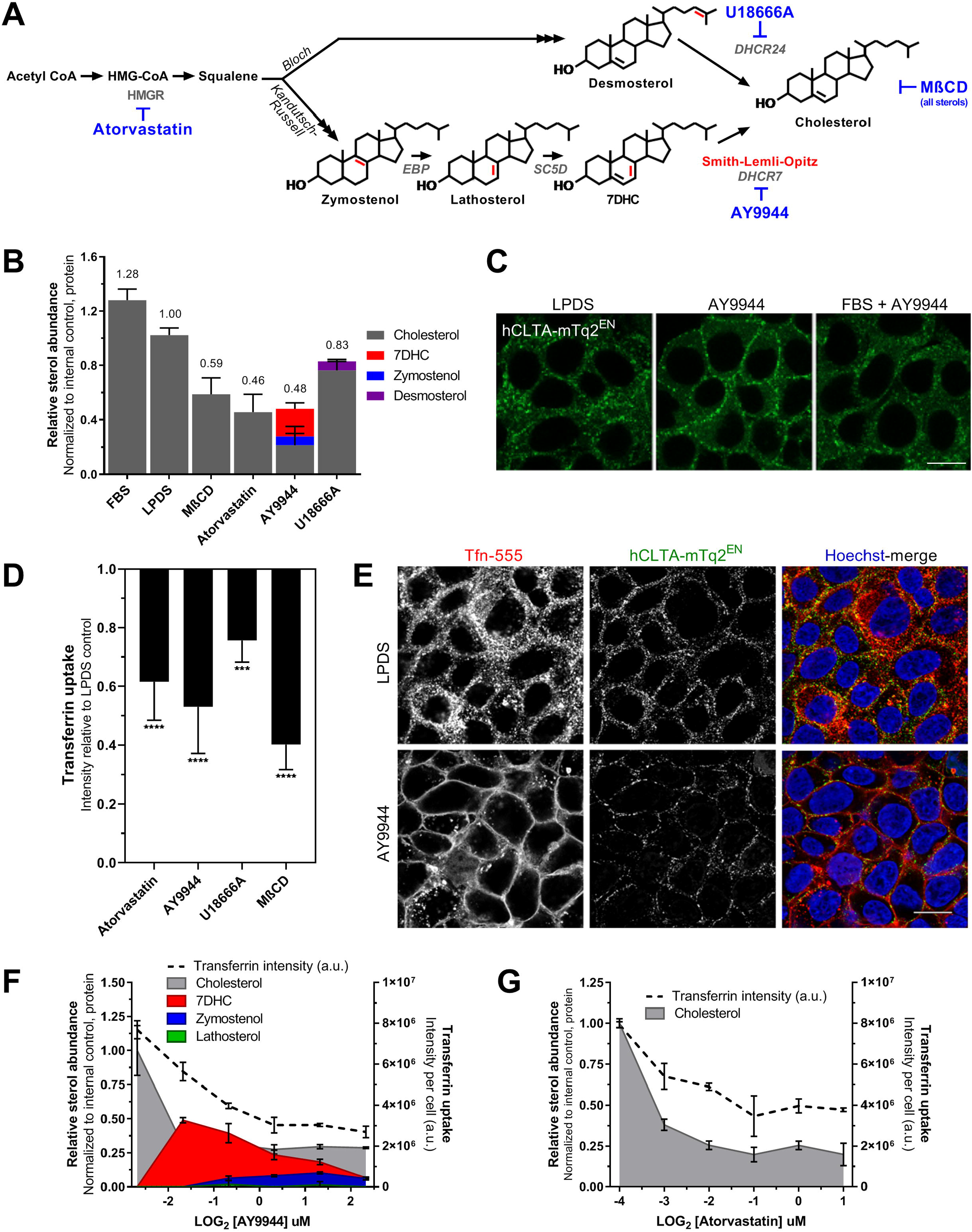
Sterol homeostasis is required for endogenous clathrin trafficking. **A)** Schematic illustrating the final steps of post-squalene cholesterol synthesis. Structural alterations of sterol precursors relative to cholesterol are indicated in red. Small molecules used in this study are noted in blue. **B)** Sterol profiles of HEK293T hCLTA^EN^-mTq2 cells cultured in cholesterol-replete FBS or 7.5% delipidated (LPDS) serum ± treatment with Atorvastatin, AY9944, or U18666A for 48 h. Acute cholesterol depletion was achieved by 1h MβCD treatment (Mean ± SD). **C)** FBS prevents AY9944-mediated CME trafficking defects. Scale bar = 10 μm. **D)** Tfn uptake under sterol depleted conditions relative to controls cultured in 7.5% LPDS for 48 h (Mean ± SD). ***, P < 0.001; ****, P < 0.0001; one-way ANOVA (F(4,40) = 38.57, p < 0.0001) and Dunnett’s test versus LPDS control (N = 9 biological replicates from 3 independent experiments, 2,000 – 4,000 cells per replicate). **E)** Sterol depletion results in accumulation of clathrin at the cell periphery and functional impairment of Tfn internalization by confocal microscopy in HEK293T hCLTA^EN^-mTq2 cells. Scale bar = 10 μm. **F, G)** Tfn uptake correlates with sterol content by GC/MS analysis (Mean ± SD). N = 3 biological replicates for Tfn uptake and N = 2 biological replicates for sterol analysis performed in parallel.

To monitor the effects of cholesterol homeostasis on CME, we used CRISPR/Cas9 gene editing to fluorescently tag clathrin light chain A (CLTA) in HEK293T using our previously described approach (**Figure S1A**) (Anderson et al., 2018; Scott et al., 2018). A pooled population of genome-edited cells was enriched by FACS to create a homogenous population of CLTA^mTq2+^ cells, thereby mitigating potential off-target effects that could arise in single cell clones (**Figure S1B, C).** Live-cell confocal microscopy taken at midsection planes revealed dynamic PM budding and rapid trafficking of clathrin-mTq2^+^ vesicles in LPDS cultured cells. All sterol depleted conditions, exemplified by AY9944 treatment, lead to striking immobilization of clathrin vesicles at the PM despite ongoing intracellular transport (**Video 1, panels 1-2**). Trafficking deficits induced by AY9944 treatment were efficiently rescued by addition of cholesterol loaded methyl-β-cyclodextrin (MβCD-Chol) as observed by rapid redistribution of clathrin-mTq2^+^ traffic from the PM, indicating that plasma membrane cholesterol alone is sufficient and essential for clathrin trafficking (**Video 1, panel 3**). This effect was rapid, restoring CME within minutes of MβCD-Chol addition and rendering mTq2^+^-clathrin traffic indistinguishable from LPDS controls within 1 h (**Video 1, panel 3**). To exclude potential secondary effects from the lipophilic properties of the cholesterol synthesis inhibitors, small molecules were administered in the presence of exogenous cholesterol, thereby minimizing endogenous synthesis (Wassif et al., 2005). Under these conditions, the cholesterol synthesis inhibitors did not produce clathrin trafficking deficits (**Figure 1C**).

To determine the extent of functional CME impairment, we used a high-content, non-biased quantitation imaging assay to analyze the internalization of fluorescently-conjugated transferrin (Tfn) across sterol depleted conditions. The degree of sterol depletion achieved with these inhibitors (**Figure 1B**), predicted the reduction in CME. Specifically, treatment with either AY9944, Atorvastatin, or MβCD reduced both sterol abundance and Tfn internalization by approximately 50%. U18666A had a smaller effect (~20% reduction) on both sterols and CME (**Figure 1D**). Confocal optical sections taken at midplane also confirmed the inhibition of Tfn internalization (**Figure 1E),** as seen by the loss of Tfn-555 positive vesicles in the cytoplasm and localization to the PM. Although Tfn internalization was impaired, high Tfn-555 signal from the PM suggests that the Tfn receptor also accumulated on the PM. Because CME was impaired regardless of the step of cholesterol synthesis inhibition or time of cholesterol depletion, we speculated that cholesterol depletion or decreased total sterol content was the major mediator of this phenomenon, rather than accumulation of specific sterol precursors.

Parallel comparison of sterol profiles by GC/MS and Tfn uptake as a function of inhibitor concentration demonstrated that CME activity was dependent on total sterol content (**Figure 1F, G)**.

Recovery of CME traffic was also predicted by sterol abundance and availability. Addition of MβCD-Chol following sterol depletion with AY9944 rapidly restored both mTq2^+^-clathrin traffic and transferrin internalization (**Figure 2B, top**). In contrast, clathrin-mTq2^+^ vesicle trafficking required 6-12 h to normalize following addition of FBS, indicating that lipoprotein-derived (LDL) cholesterol uptake and redistribution was much slower (**Figure 2B, middle**), likely due to depressed CME-dependent LDL uptake (Davis et al., 1986). Clathrin-mTq2^+^ vesicle trafficking also did not recover upon removal of AY9944 and addition of LPDS within 12 h, indicating the rate of CME rescue by endogenous cholesterol synthesis is much slower than either direct or LDL-dependent cholesterol uptake (**Figure 2B, bottom**). Feedback inhibition of 7DHC on the synthetic pathway may also limit sterol biosynthesis (Honda et al., 1998). Cellular sterol profiles corresponded with the resumption of CME (**Figure 2C**). Notably, incubation of AY9944 depleted cells with MβCD-Chol corrected cellular sterol profiles more completely than availability of LDL-derived cholesteryl esters over a 12 h period (**Figure 2C**), suggesting greater efficacy and bioavailability of cholesterol supplementation independent of endocytic traffic. These rescue assays demonstrate a definitive requirement for an optimal concentration of cellular cholesterol to facilitate clathrin vesicle trafficking.

**Figure 2.**
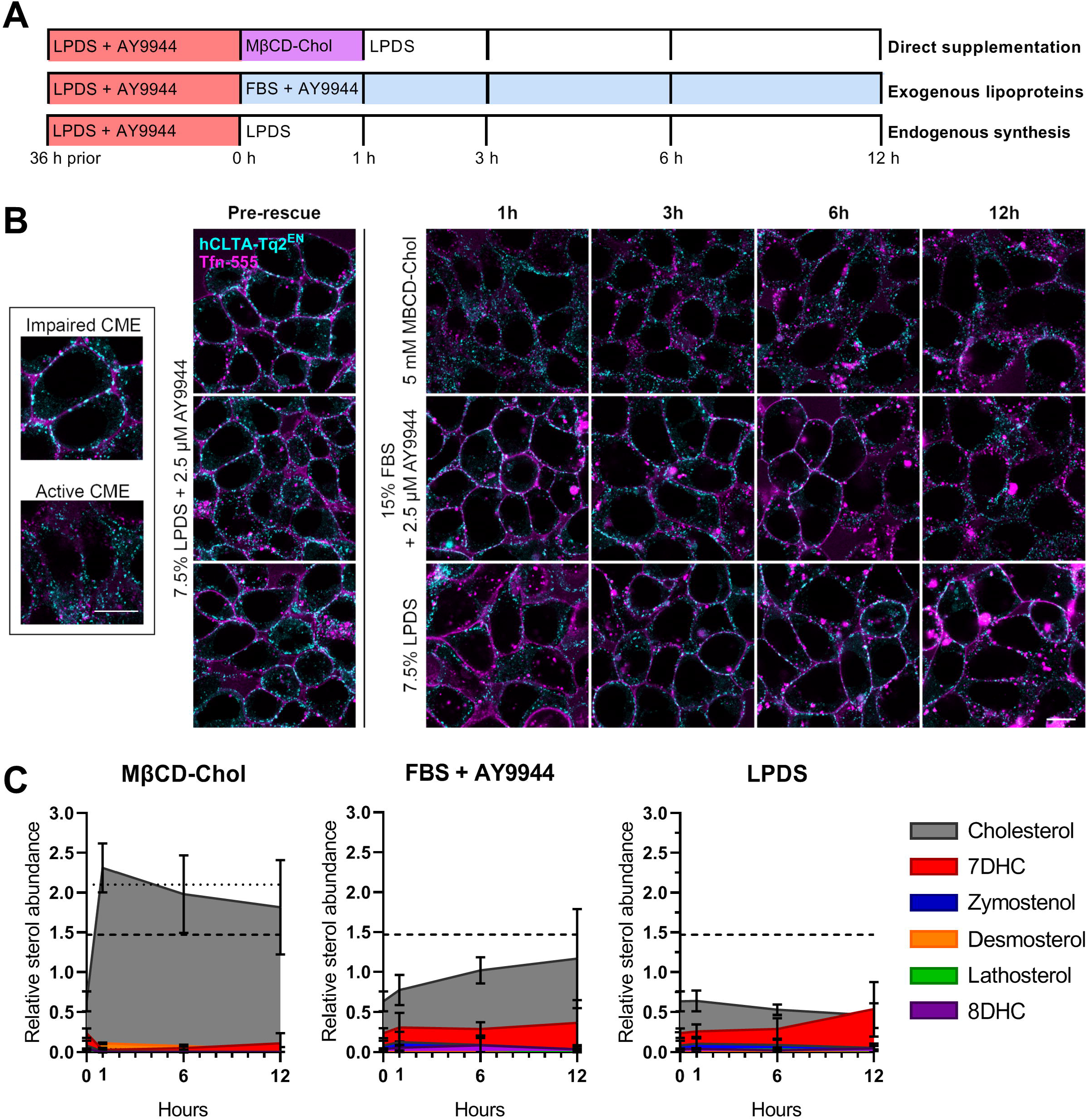
AY9944 induced clathrin trafficking deficits are rescued by direct cholesterol loading with MβCD. **A)** Overview of recovery conditions. Following AY9944 sterol depletion, HEK293T hCLTA-Tq2^EN^ cells were exposed to cholesterol-loaded MβCD (MβCD-Chol, and chased in LPDS media) **[Top]**, supplemented continuously with lipoprotein-rich 15% FBS in the presence of AY9944 **[Middle]**, or incubated continuously in 7.5% LPDS media in the absence of AY9944 to allow endogenous cholesterol synthesis **[Bottom]**. **B)** Mid-plane live-cell confocal microscopy of HEK293T hCLTA-Tq2^EN^ following addition of transferrin-conjugated to AF-555 (Tfn-555) during recovery as described in (A). Representative images from different fields of view shown. Tfn and clathrin distributions in normal and arrested CME **[Inset]**. Scale bar = 10 μm. **C)** Corresponding cellular sterol profiles quantified by GC/MS (Mean ± SD). N = 3 biological replicates from independent experiments. Dotted and dashed lines represent total sterol abundance of untreated FBS and LPDS cultured cells, respectively.

### 7-dehydrocholesterol can compensate for cholesterol in facilitating CME

As SLOS leads to a significant accumulation of 7DHC (Irons et al., 1993; Tint et al., 1994), we wanted to directly address the ability of 7DHC to compensate for the activity of cholesterol with regards to supporting CME. To this aim, rescue of aberrant clathrin trafficking following AY9944 treatment was evaluated by loading of cholesterol or 7DHC directly to cellular membranes using MβCD as a water-soluble carrier. Incubation with MβCD-7DHC efficiently rescued total sterol levels back to cholesterol-replete conditions with 7DHC being the predominate form (~80% of total sterols) (**Figure S2A**). Addition of uncomplexed MβCD to cells already depleted of sterols did not appreciably further reduce sterol levels, which may reflect diminished “free” cholesterol available for MβCD exchange. Restoration of total sterol levels by either MβCD-Chol or MβCD-7DHC resulted in resumed clathrin-mTq2^+^ trafficking (**Video 2**) and robust Tfn internalization (**Figure S2B, C**). This finding suggests that loss of planarity within the sterol ring, due to the additional double bond in 7DHC (Serfis et al., 2001), does not impede its endocytic function at the membrane. Although 7DHC has been suggested to impact other aspects of PM structure including bending rigidity (Gondre-Lewis et al., 2006) and lipid packing (Tulenko et al., 2006), our results demonstrate 7DHC can compensate for cholesterol in sustaining CME activity. Thus, CME appears to be sensitive to total sterol content but robust to sterol identity.

### Altered sterol homeostasis inhibits clathrin-coated pit dynamics and membrane bending

Cholesterol’s multifaceted influence on PM properties likely contributes to lowering the energetic cost of membrane remodeling (Bruckner et al., 2009). Because EM studies following MβCD treatment indicate arrest at preliminary stages of pit bending (Rodal et al., 1999; Subtil et al., 1999), sterol depletion may inhibit membrane curvature generation and recruitment of CME proteins. To determine how cholesterol and sterols influence membrane bending during clathrin assembly, we used polarized total internal reflection fluorescence microscopy (polTIRFM) of DiI, which allows imaging of PM topography (Aguet et al., 2013; Anantharam et al., 2010; Scott et al., 2018). Imaging of membrane bending in polTIRFM is achieved by selective excitation of DiI-labeled PM within the vertical (P polarization) or horizontal (S polarization) plane with respect to the coverslip and quantified by the ratio of P/S fluorescence (Anantharam et al., 2010). We previously demonstrated that this approach can accurately measure changes in membrane topography during membrane bending during CME by correlative light and electron microscopy (CLEM) and atomic force microscopy (CLAFM) (Scott et al., 2018). The membrane bending dynamics of individual endocytic events were recorded in SK-MEL-2 cells endogenously expressing clathrin-mTq2^+^ and dynamin-eGFP^+^ with DiI labeled membranes (**Video 3**). CME dynamics of the SK-MEL-2 cell line have been well characterized by live-cell TIRFM and polTIRFM (Doyon et al., 2011; Scott et al., 2018).

Upon disruption of cholesterol homeostasis, total sterol content relative to LPDS culture was found to be 147 ± 24% (FBS), 81 ± 7% (Atorvastatin), 82 ± 11% (AY9944), or 34 ± 9% (MβCD) in SK-MEL-2 treated cells (data not shown). Under all sterol deplete conditions (AY9944, Atorvastatin, and MβCD), the total number of unique, distinct clathrin assembly spots per cell area over time was significantly lower relative to cells grown in complete media or LPDS (**Figure 3C**). Both moderate and severe sterol depletion led to an increase in the number of persistent clathrin tracks (>400 s) (**Figure 3D**) and average clathrin lifetimes per cell (**Figure 3F**). Accumulation of clathrin-mTq2^+^ spots with sterol depletion indicated that the reduction in clathrin tracks was not due to lack of clathrin assembly, but arrested CME (**Video 4**, **Figure 3A**). Clathrin-mTq2^+^ tracks were further categorized according to detection of dynamin and P/S signal. Although dynamin may contribute to early stages of clathrin-coated pit maturation (Aguet et al., 2013; Srinivasan et al., 2018), it is most abundant during membrane fission. Dynamin-eGFP^+^ was visualized as a burst of accumulation immediately prior to the completion of endocytosis and the disappearance of the clathrin-coated vesicle (**Figure 3B**). Using dynamin recruitment as a marker of completed CME events, we found that the number of productive CME events per cell area was dramatically reduced during sterol depletion (**Figure 3E**). When CME events did occur, they exhibited longer lifetimes compared with FBS or LPDS controls (**Figure 3G, Figure S3**). Following AY9944 treatment, the average lifespan of CME events which could be observed was increased by 40 s (95% CI [89,148]) compared to LPDS controls (95% CI [66, 97]). Moreover, CME lifetimes in AY9944 treated cells are underestimated by this data due to temporal limitations in video tracking of persistent CME tracks. A detailed breakdown of clathrin lifetimes under control or sterol depleted conditions is shown in **Figure S4**.

**Figure 3.**
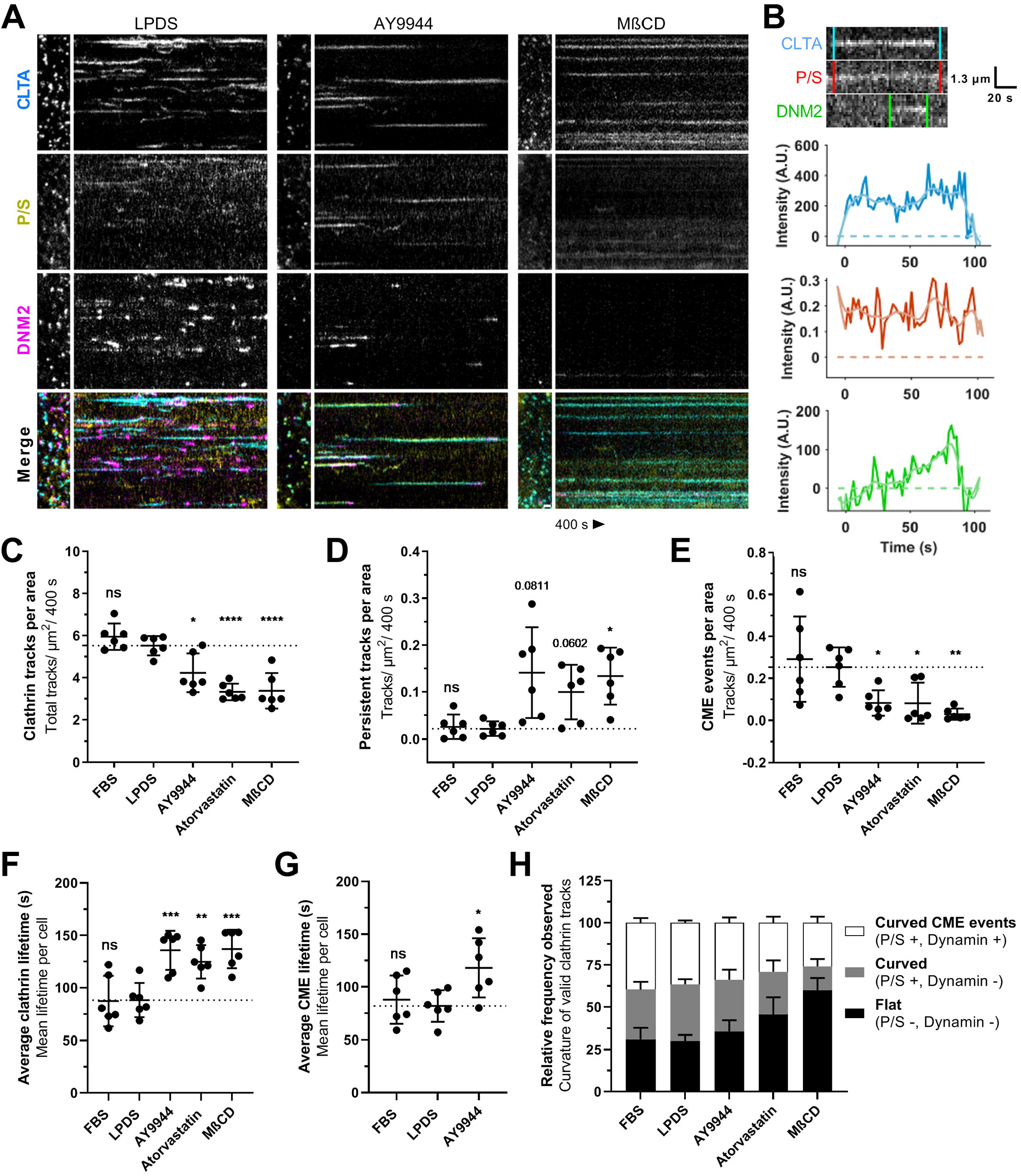
Loss of cholesterol homeostasis disrupts clathrin-coated pit dynamics. **A)** Representative polTIRFM kymographs comparing SK-MEL-2 (hCLTA-mTq2^EN^/hDNM2-eGFP^EN^) cells grown in 10% LPDS **[Left]** to moderate sterol depletion with addition of 2.5 μM AY9944 **[Middle]** or severe sterol depletion with 5 mM MβCD **[Right]**. Positive curvature generation is indicated by the ratio of P/S fluorescence in DiI labeled membrane. Time = 400 s. Scale bar = 1μm. **B)** Representative kymograph of CLTA-mTq2, membrane bending (P/S), and DNM2-eGFP and corresponding intensity traces during a single CME event. **C)** Unique clathrin events tracked per cell area analyzed across a 400 s time-lapse acquisition (Mean ± SD). *, P < 0.05; ****, P < 0.0001; one-way ANOVA (F(4,25) = 18.86, p < 0.0001) and Dunnett’s test versus LPDS control (N = 6 cells from 2-4 independent experiments, 3,000-30,000 tracks analyzed per cell). **D)** Number of persistent tracks (lifetime > 400 s) per cell area analyzed (Mean ± SD). Adjusted P values; *, P < 0.05; Welch ANOVA (F(4, 11.43) = 7.531, p = 0.0032) and Dunnett’s T3 test versus LPDS control (N = 6 cells from 2-4 independent experiments). **E)** Productive CME events per cell area (Mean ± SD). *, P < 0.05; **, P < 0.01; Welch ANOVA (F(4,11.24) = 8.948, p = 0.0017) and Dunnett’s T3 test versus LPDS control (N = 6 cells from 2-4 independent experiments). **F)** Average clathrin lifetimes per cell across the indicated treatment groups (Mean ± SD). Lifetime analysis was limited by truncation of clathrin tracks extending beyond the first or last frame (400 s). **, P < 0.01; ***, P < 0.001; one-way ANOVA (F(4,25) = 10.46, p < 0.0001) and Dunnett’s test versus LPDS control (N= 6 cells from 2-4 independent experiments). **G)** Average lifetime of observable CME events per cell across the indicated treatment groups (Mean ± SD). Lifetime of CME limited to observed events > 400 s, such that true lifetimes exceed the estimates presented (400 s). *, P < 0.05; one-way ANOVA (F(4,15) = 4.408, p < 0.0312) and Dunnett’s test versus LPDS control (N= 6 cells from 2-4 independent experiments). **H)** Relative frequency of clathrin tracks observed associated with curvature generation (Mean ± SEM, N = 6 cells). Flat clathrin tracks were classified as exhibiting neither curvature nor dynamin recruitment, may represent persistent tracking or abortive clathrin events visiting the TIRM field. Curved clathrin events were classified as exhibiting positive P/S curvature signal, but failed to recruit dynamin. Curved CME events were classified by the ability of clathrin positive events to recruit dynamin.

While both moderate (AY9944 and Atorvastatin) and severe (MβCD) cholesterol depletion led to diminished rates of CME, we observed treatment-specific impacts on membrane curvature dynamics. Cells displaying ~20% reduction in total sterol content following AY9944 or Atorvastatin treatment exhibited clathrin-mTq2^+^ spots associated with strong P/S signals, indicating the presence of highly curved pits which stalled prior to acquiring bursts of dynamin (**Figure 3A**). DMN2-eGFP^+^ localization was delayed in both AY9944 and Atorvastatin treated cells compared to LPDS controls, suggesting clathrin events inefficiently recruited dynamin and many did not become productive CME events. As illustrated by kymographs of long-lived clathrin tracks failing to undergo scission (**Figure S5**), fluctuations in P/S membrane bending along clathrin tracks were noted. While weak DNM2^eGFP+^ signal was associated with many curved clathrin structures, robust dynamin signatures could also be seen dissociating without a loss of clathrin signal. No differences in dynamin lifetimes were observed in association with productive CME events (data not shown). These findings suggest moderate disruption of cholesterol synthesis allowed membrane bending during the early stages of CME, but precluded neck formation and the recruitment of dynamin. In contrast, cells displaying ~65% reduction in total sterols following MβCD treatment exhibited diminished P/S signals with little to no dynamin recruitment (**Figure 3A, H**), indicating that severe sterol depletion inhibits CME at the stage of initial membrane bending. This result is consistent with previous studies describing flat clathrin-coated structures following MβCD mediated sterol depletion, consistent with an inability of clathrin-coated pits to invaginate (Rodal et al., 1999). Altogether, these findings suggest a homeostatic level of cholesterol is required at multiple stages of PM bending for the CME pathway to proceed normally.

### Sterol depletion inhibits curvature generation and neck formation during CME, producing asymmetric pit structure

In order to understand the effects of sterol depletion on membrane curvature during CME, we preformed platinum replica EM of unroofed SK-MEL-2 cells to quantitatively assess ultrastructure (Sochacki et al., 2017; Sochacki et al., 2014). We classified clathrin structures as flat, shallow, domed, or spherical to quantitatively assess curvature status (**Figure 4B**). In cholesterol replete conditions (FBS or LPDS culture), flat and spherical clathrin-coated pits were most commonly observed with fewer shallow or domed structures. This observation was consistent with rapid progression of CME pits from flat to spherical as previously described in SK-MEL-2 cells cultured in FBS (Scott et al., 2018). In contrast, moderate sterol depletion with either Atorvastatin or AY9944 increased the relative frequency of shallow and domed structures, predominately at the expense of spherical pits (**Figure 4B**). The increased frequency of shallow/domed structures explains the high P/S signal observed on clathrin spots by polTIRFM with AY9944 and Atorvastatin treatment (**Figure 3A**). Combined with the fluctuations observed in the P/S signals (**Figure S5**), these findings suggest that the structural transition from shallow to domed morphology is metastable, potentially due to the energetics of clathrin-coat lattice reorganization competing with elevated free energy costs of membrane bending upon sterol depletion. Consistent with polTIRFM studies, severe sterol depletion by MβCD shifted the relative frequency of coated pit curvature to predominately flat or shallow coated pits, reinforcing the concept that sterol depletion from the PM dramatically increases the energetic cost of membrane bending (**Figure 4A**). Absolute values of the categorized clathrin structures per cell are summarized in **Figure S6**. In support of intrinsic membrane properties altering the energetics of membrane bending, caveolae under cholesterol deplete conditions also exhibited a flattened, disassembled morphology (**Figure 4F**) and were decreased in number at the plasma membrane (**Figure 4G**). These findings provide direct structural evidence that sterol abundance influences the progression of CME events by altering membrane bending dynamics at multiple stages of curvature generation.

**Figure 4.**
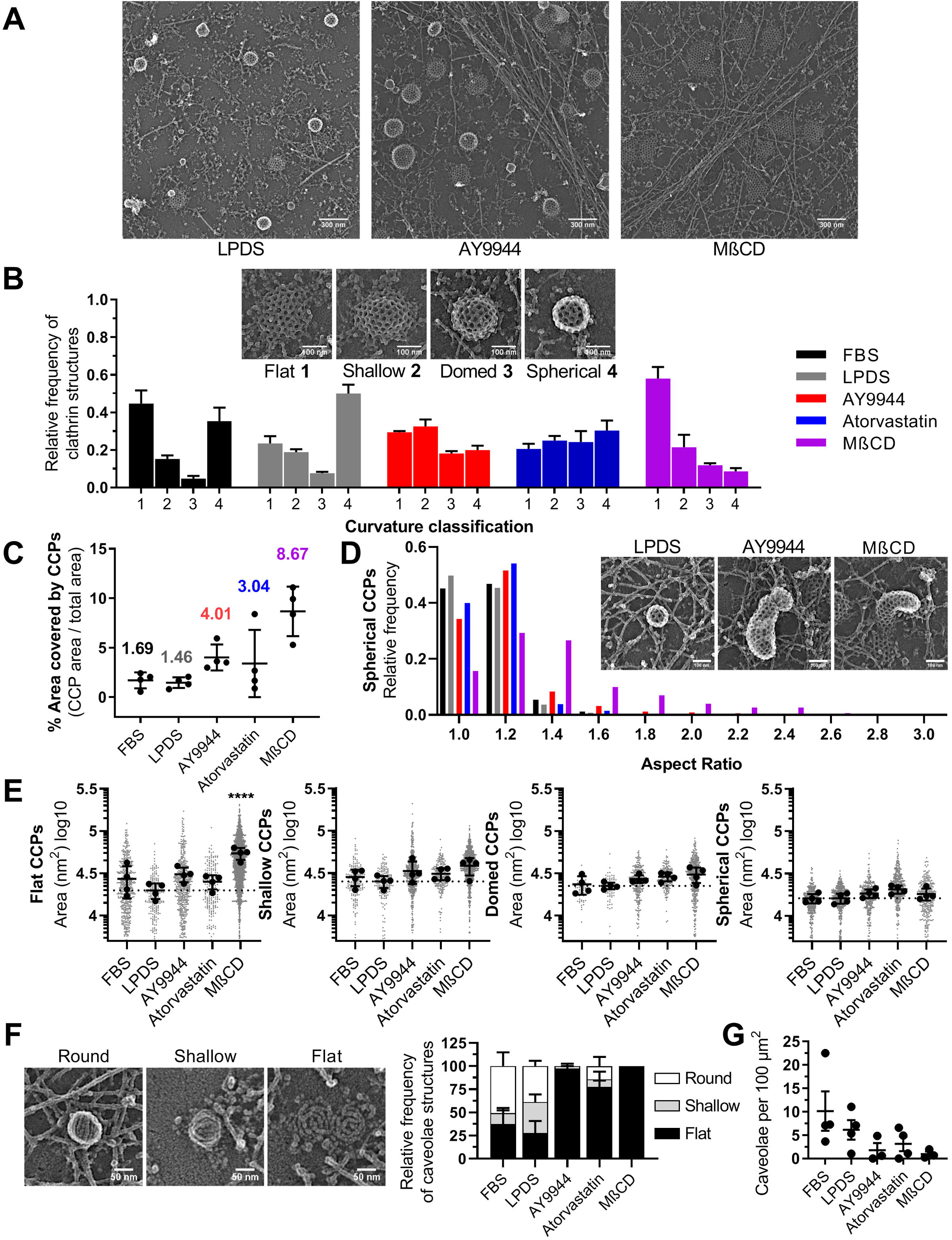
Aberrant clathrin-coated pit ultrastructure under conditions of sterol depletion. **A)** Representative platinum-replica TEM images from LPDS cultured cells, AY9944 treated cells, and MβCD treated cells. **B)** Relative frequency of clathrin structures by curvature classification (Mean ± SEM). **C)** Area of unroofed cells covered by clathrin structures (Mean ± SD). **D)** Histogram of spherical clathrin structures showing the distribution of coated pit asymmetry by aspect ratio (major/minor width). **E)** Average area of clathrin-coated structures by curvature classification relative to LPDS controls (Mean ± SD). ****, P < 0.0001; one-way ANOVA (F(4,15) = 12.91, p < 0.0001) and Dunnett’s test versus LPDS control. N = 4 replicas per condition from two independent experiments. Total number of clathrin structures analyzed: FBS = 1164; LPDS = 988; AY9944 = 1700; Atorvastatin = 1003; MβCD = 3818. **F)** Representative platinum-replica TEM images of caveolae ultrastructure and relative frequency of observed structures by curvature classification (Mean ± SEM). **G)** Total number of caveolae structures on the membrane per area (Mean ± SEM). N = 3-4 replicas per condition from two independent experiments. Total number of caveolae structures analyzed: FBS = 171; LPDS = 89; AY9944 = 21; Atorvastatin = 31; MβCD = 13.

Sterol depletion also resulted in loss of structural symmetry within pits. While most clathrin structures were flat in MβCD treated cells, the few observable spherical pits were asymmetric and distorted (**Figure 4D**), suggesting aberrant energetics of membrane bending. Upon sterol depletion with both AY9944 and Atorvastatin, distorted coated pits exhibiting elongated necks without a narrow base were also observed (**Figure 4D inset**) and the quantified asymmetry scaled with the severity of cholesterol depletion such that MβCD>>AY9944~Atorvastatin (**Figure 4D)**. Aside from the numerous flat clathrin structures within MβCD treatment which were beginning to converge (**Figure 4E, panel 1**), no significant differences in the average size of clathrin structures were observed across treatments within the shallow, domed, and spherical subsets of curvature classification (**Figure 4E, panels 2-4**). This suggests that once energetic barriers were crossed, general pit dimensions were constant. Notably, the area of the PM covered by clathrin increased dramatically from approximately 1.5% in FBS and LPDS controls to 3.0%, 4.0%, and 8.7% when treated with Atorvastatin, AY9944, or MβCD, respectively (**Figure 4C**). A possible explanation for the loss of clathrin pit symmetry upon sterol depletion may be due to the accumulation of stalled pits on the PM resulting in asymmetric membrane bending trajectories during the remodeling of large flat clathrin lattices into vesicles (Avinoam et al., 2015; Scott et al., 2018). This data demonstrates that perturbation of cholesterol homeostasis alters the process of clathrin assembly and the trajectory of membrane bending leading to an accumulation of normal and distorted clathrin-coated pits on the PM.

### Phase separation predicts the sterol structural requirements for supporting CME

Considering that membrane bending was impacted at both early and late stages of pit formation congruent upon total sterol abundance, the ability of a structurally dissimilar sterol (7DHC) to support CME, and the relative abundance of cholesterol within the PM, we wanted to further define the biophysical properties of sterols necessary for facilitating curvature and CME. Biophysical studies in model membranes have suggested rapid transbilayer redistribution (flip-flop) of cholesterol relieves stress associated with membrane remodeling by alleviating the molecular disparities between the inner and outer PM leaflets during bending (Bruckner et al., 2009; Lange et al., 1981). In contrast to other lipid components of the PM, cholesterol transbilayer flip-flop occurs within milliseconds (Choubey et al., 2013; Hamilton, 2003; Steck et al., 2002), a timescale sufficiently fast to support local membrane remodeling during an endocytic event, which typically takes 80-100 s in the case of CME (Taylor et al., 2011). Additionally, cholesterol and select sterols have the capacity to partition bilayers into liquid-ordered (*L_o_*) or liquid-disordered (*L_d_*) phases due to preferential tight packing of sterols and lipid components. Phase separation is postulated to underlie membrane shape transitions by minimizing line tension between domain boundaries (e.g. driving progressive curvature generation to neck structures) (Jülicher and Lipowsky, 1993, 1996) and is implicated in membrane curvature and budding behavior in artificial membranes (Bacia et al., 2005).

To clarify the sterol structural requirements necessary to support CME, we used MβCD to deliver structurally diverse sterols and tested their ability to rescue CME function following AY9944 inhibition. As summarized in **Figure 5A, Table S2**, sterols were chosen based on differences in ring structure (lathosterol, 7DHC), aliphatic tail structure (desmosterol), and predicted ability to phase separate or support transbilayer movement of sterols. Cholestenone, which substitutes the 3β-hydroxyl with a ketone group as typical of steroid hormones, likely diminishes polar interactions at the membrane-water interface to promote more rapid interleaflet exchange (Róg et al., 2008). Conversely, increased polarity (4β-hydroxycholesterol) or charge (cholesterol sulfate) would impede sterol exchange between membrane leaflets and do not support phase separation (Bacia et al., 2005; Wenz and Barrantes, 2003). Incubation with desmosterol, 7DHC, lathosterol or 4β-hydroxycholesterol MβCD-sterol complexes efficiently rescued total sterol levels to LPDS levels (**Figure 5B**). MβCD-cholesterol sulfate and cholestenone complexes only allowed for sterol exchange (total cellular sterol content remained near AY9944 levels). Sterols previously demonstrated to allow for sterol flipping and supporting phase separation (desmosterol, 7DHC) rescued Tfn internalization (**Figure 5C, D**). However, Tfn internalization was not rescued by sterols which do not support phase separation or allow membrane flip-flop (4β-hydroxycholesterol, galactosyl cholesterol) (**Figure 5C, D**). This suggests that sterol structures supporting phase separation confer similar requirements necessary to support CME.

**Figure 5.**
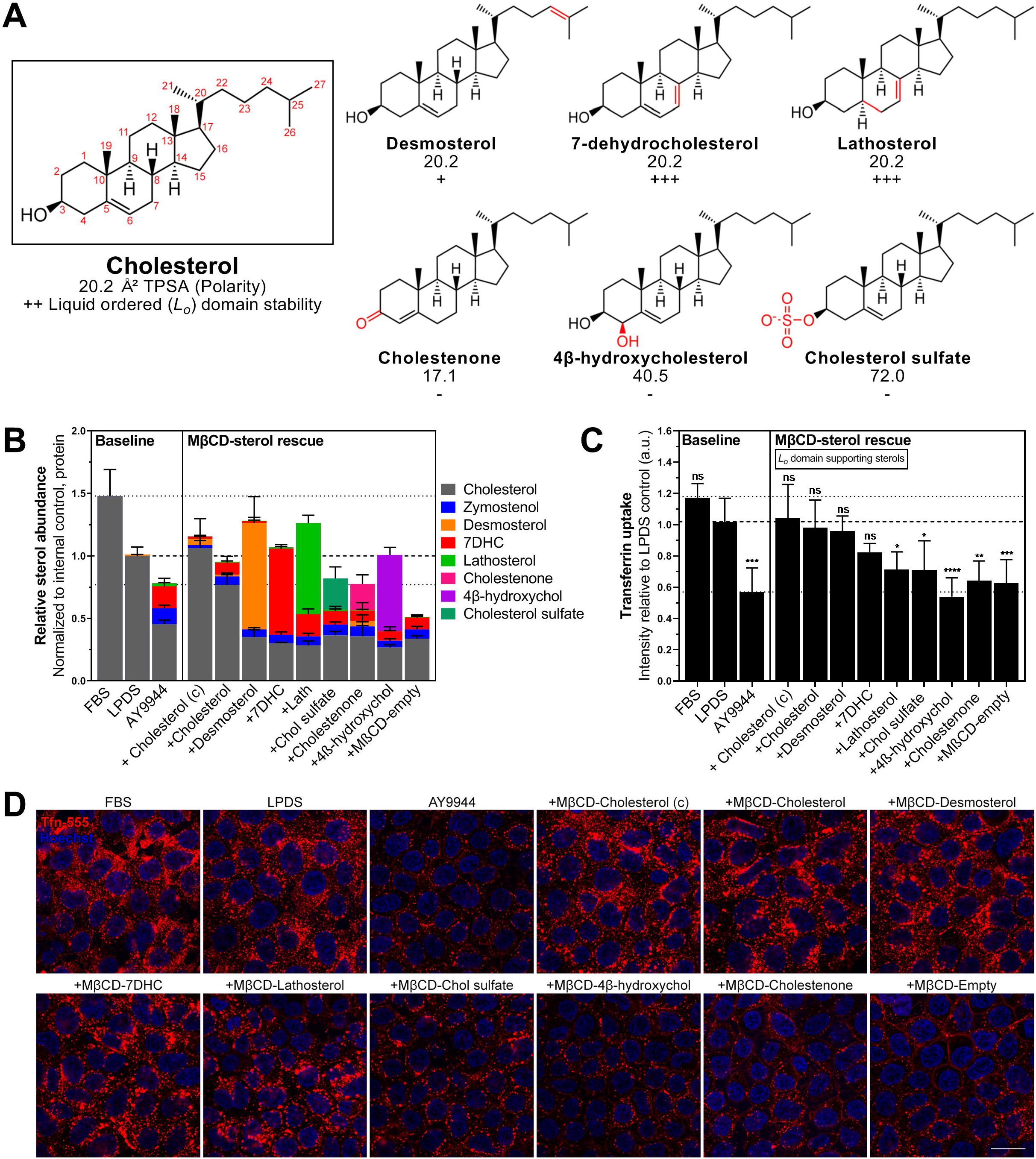
Sterol mediated phase separation is required to support CME. The ability of structurally diverse sterols to rescue CME inhibition due to sterol depletion by AY9944 treatment was evaluated by direct delivery of sterols to the PM via MβCD carrier. **A)** Summary of physical properties and phase separation behavior of sterols. Topological Polar Surface Area (TPSA) computed surface sum over all polar atoms. Ordered lipid (*L_o_*) domain (raft) stabilizing (+) or disrupting (−) sterols indicated relative to cholesterol (++). Refer to **Table S2** for additional details. **B)** Sterol profiles of AY9944 treated HEK293T cells following 1h incubation with sterol loaded MβCD (Mean ± SD). N = 4 independent biological replicates from two MβCD-sterol preparations. **C)** Tfn uptake relative to controls cultured in 7.5% LPDS for 48 h (Mean ± SD). *, P < 0.05; **, P < 0.01; ***, P < 0.001; ****, P < 0.0001; one-way ANOVA (F(12, 51) = 11.23, p < 0.0001) and Dunnett’s test versus LPDS control (N = 5 biological replicates from 3 independent experiments, ~1,500 cells per replicate). Cholesterol, desmosterol, 7DHC, and lathosterol support formation of *L_o_* domains in model membranes as designated. **D)** Representative confocal images taken mid-plane following 30 min incubation with AF-555 conjugated transferrin (Tfn). Scale bar = 20 μm. (c), Commercially available, pre-loaded MβCD-Chol.

### Integrity of the actin cytoskeleton does not significantly modulate CME sterol dependence

During CME, the polymerization energy of the clathrin lattice is opposed by membrane bending energy (Saleem et al., 2015). While the actin cytoskeleton is not absolutely required for CME in mammalian cells, it becomes increasingly important as membrane tension increases (Batchelder and Yarar, 2010; Boulant et al., 2011). Actin polymerization is thought to provide additional force during the transition of domed to spherical shaped pits in preparation for the vesicle scission phase (Grassart et al., 2014), (Hassinger et al., 2017; Skruzny et al., 2012). Inhibition of CME dynamics within mitotic cells has been observed where rigid cortical actin and increased membrane tension dominate (Kaur et al., 2014). Moreover, cytoskeletal alterations which occur following cholesterol depletion may enhance membrane-cytoskeletal adhesions, resulting in either sequestration of actin away from CME or elevated membrane tension (Ayee and Levitan, 2016; Khatibzadeh et al., 2012). To determine whether inhibition of CME by sterol depletion is mediated through changes to the actin cytoskeleton or specific to the membrane, we depolymerized actin with latrunculin B pre-treatment during inhibition by AY9944 alone or following recovery of CME activity by sterol repletion (**Figure S7C**). We did not find evidence of significant latrunculin B sensitivity in either restoring CME activity in sterol depleted cells or inhibiting MβCD-cholesterol rescue (**Figure S7E**). These data suggest the influence of sterols on CME is largely independent of actin remodeling but directly dependent on changes to the physical properties of the PM.

### Patient fibroblasts containing mutations within cholesterol synthesis genes exhibit functional CME deficits

To determine if CME activity was reduced within the context of a human disease characterized by sterol disruption, Tfn uptake assays were performed on fibroblasts derived from SLOS subjects. Three cell lines were selected to be representative of the SLOS phenotypic spectrum (**Table S3**). Compared to an unaffected control, Tfn uptake was greatly reduced in SLOS samples upon culture within LPDS conditions (**Figure 6A, D**), responding similarly to AY9944 treated control fibroblasts. While CME activity was similarly inhibited in all three SLOS fibroblasts when cultured in LPDS conditions, CME function correlated to the total sterol levels of each cell line as determined by GC/MS (**Figure 6B**). Notably, even under cholesterol replete conditions, fibroblasts from the most biochemically severe SLOS cell lines trended towards diminished Tfn internalization, suggesting that exogenous lipid supplementation and lipoprotein is unable to fully rescue CME function (**Figure 6C**). These data demonstrate a direct link between sterol homeostasis and CME within SLOS subjects, while suggesting functional deficits in CME may have clinical impact within this patient population.

**Figure 6.**
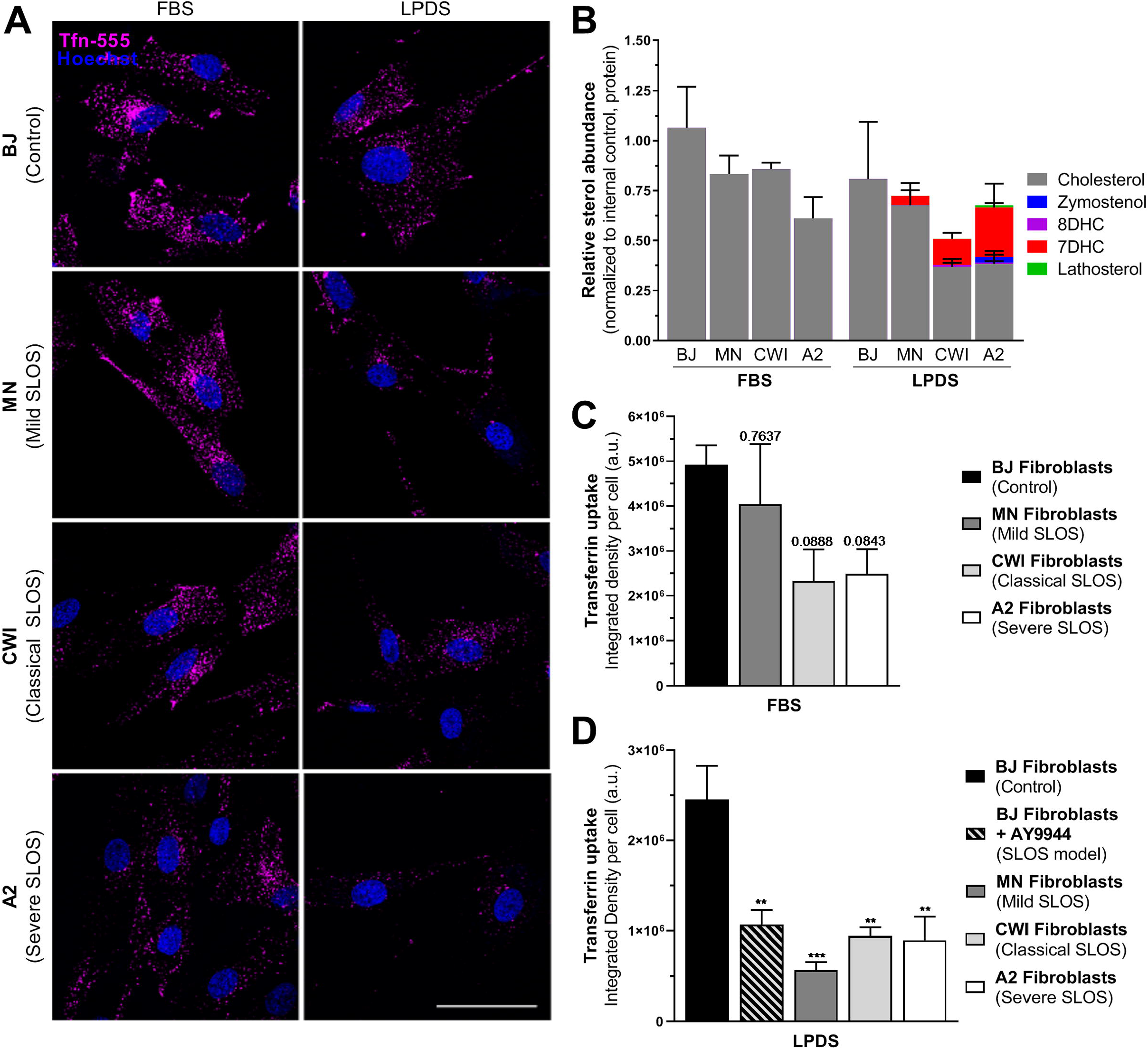
CME dynamics are inhibited within Smith-Lemli-Opitz syndrome patient-derived fibroblasts. **A)** Representative images demonstrating AF-555 conjugated Tfn uptake in control and SLOS patient-derived fibroblasts grown under cholesterol replete and deplete conditions. Scale bar 50 μm. **B)** Validation of SLOS fibroblast biochemistry by GC/MS analysis. **C)** Tfn uptake relative to unaffected BJ fibroblasts grown in 15% FBS for 10 d (Mean ± SEM). Adjusted P values, one-way ANOVA (F(3,10) = 2.916, p = 0.0869) and Dunnett’s test versus BJ control (N = 3-4 independent replicates, 25 cells per replicate). **D)** Tfn uptake relative to unaffected BJ fibroblasts grown in 7.5% LPDS for 10 d (Mean ± SEM). **, P < 0.01; ***,P < 0.001, one-way ANOVA (F(4,13) = 9.464, p = 0.0008) and Dunnett’s test versus BJ control (N = 2-4 independent replicates, 25 cells per replicate).

## DISCUSSION

Here, we present evidence for the role of sterols in aiding membrane curvature generation during CME by direct observation of alterations in membrane-bending dynamics. Ultrastructure analysis and polTIRFM of clathrin-coated pits under increasing degrees of cholesterol depletion reveal domed and shallow structures respectively, supporting the model that redistribution of sterols impacts the energetics of membrane bending during both initiation of curvature and formation of the endocytic neck. We demonstrate that total sterol levels correlate with CME productivity in multiple cell types, consider the structural requirements of sterols to support CME, and find functional CME impairment in Smith-Lemli-Opitz patient samples, exhibiting *DHCR7* mutations and reduced sterol levels.

As the single most abundant lipid within the PM, the presence of cholesterol is critical to the biophysical properties of the bilayer. Our findings suggest that cholesterol is relevant to multiple stages of PM bending during an endocytic event, consistent with a structural role in alleviating energetic stress associated with curvature formation. Rescue of trafficking deficits using sterol substitution strategies suggest that the structural requirements for CME mirror the sterol requirements for supporting *L_o_* domain formation, or minimally lipid packing capacity (Megha et al., 2006; Wenz and Barrantes, 2003). This is in agreement with a report by Kim, et.al. which concludes that the presence of the 3β-hydroxyl and ability of a sterol to support ordered domain formation is necessary for endocytosis (Kim et al., 2017). In our hands, desmosterol behaved most similarly to cholesterol while the strongly raft promoting sterols, 7DHC and lathosterol, compensated for cholesterol to a lesser degree. While this observation does not imply stable PM rafts nor exclude regulation of membrane protein effectors, it is notable given a large body of work implicating membrane phase separation within curved membranes (Bacia et al., 2005; Bruckner et al., 2009; Hilgemann et al., 2020; Huttner and Zimmerberg, 2001; Jülicher and Lipowsky, 1993, 1996; Pandit et al., 2004). Budding theory originally proposed by Julicher and Lipowsky (Jülicher and Lipowsky, 1993, 1996) predicts that the boundaries between *L_o_* and *L_d_* domains tend to be minimized by “line tension”, such that induction of curvature progresses to budding of either domain. This behavior has recently been reviewed as a driving force in other clathrin-independent endocytic pathways (Hilgemann et al., 2020) and during CME (Frey and Schwarz, 2020). Immediate stress relaxation due to fast sterol transbilayer flip-flop is an underlying assumption within this model (Jülicher and Lipowsky, 1996) and, moreover, rapid sterol flip-flop explains how sterol structure determines curvature and budding phenomena within GUVs undergoing osmotic deflation (Bacia et al., 2005). During CME progression, a high energetic cost is associated with the initiation of membrane bending and is problematic for vesicle neck formation (Lentz et al., 2002). *In vitro* studies in lipid monolayers demonstrate that lipid distribution becomes non-uniform following bending with cholesterol concentrating to regions of high curvature (Wang et al., 2007). Interestingly, the effects of cholesterol on membrane rigidity (Henriksen et al., 2004; Song and Waugh, 1993) would appear contradictory to its observed effect in facilitating membrane curvature. Biophysical studies in artificial membranes suggest cholesterol flip-flop may relax tension in *L*_o_ domains despite their increased stiffness (Bacia et al., 2005; Bruckner et al., 2009), explaining why lipid rafts may be preferential sites for membrane budding and endocytosis (Conner and Schmid, 2003; Huttner and Zimmerberg, 2001). Our findings suggest that sterol-mediated stress relaxation behavior previously described within model membranes (Bacia et al., 2005; Bruckner et al., 2009) may translate to membrane remodeling *in vivo*. While it’s possible the relationship between sterol abundance and PM curvature involves a previously unrecognized and complex lipid-sensing mechanism, it seems parsimonious to consider that cholesterol passively facilitates membrane remodeling through rapid transbilayer flip-flop, relieving membrane stress between the inner and outer PM leaflets during pit formation to support high curvature (**Figure 7**).

**Figure 7.**
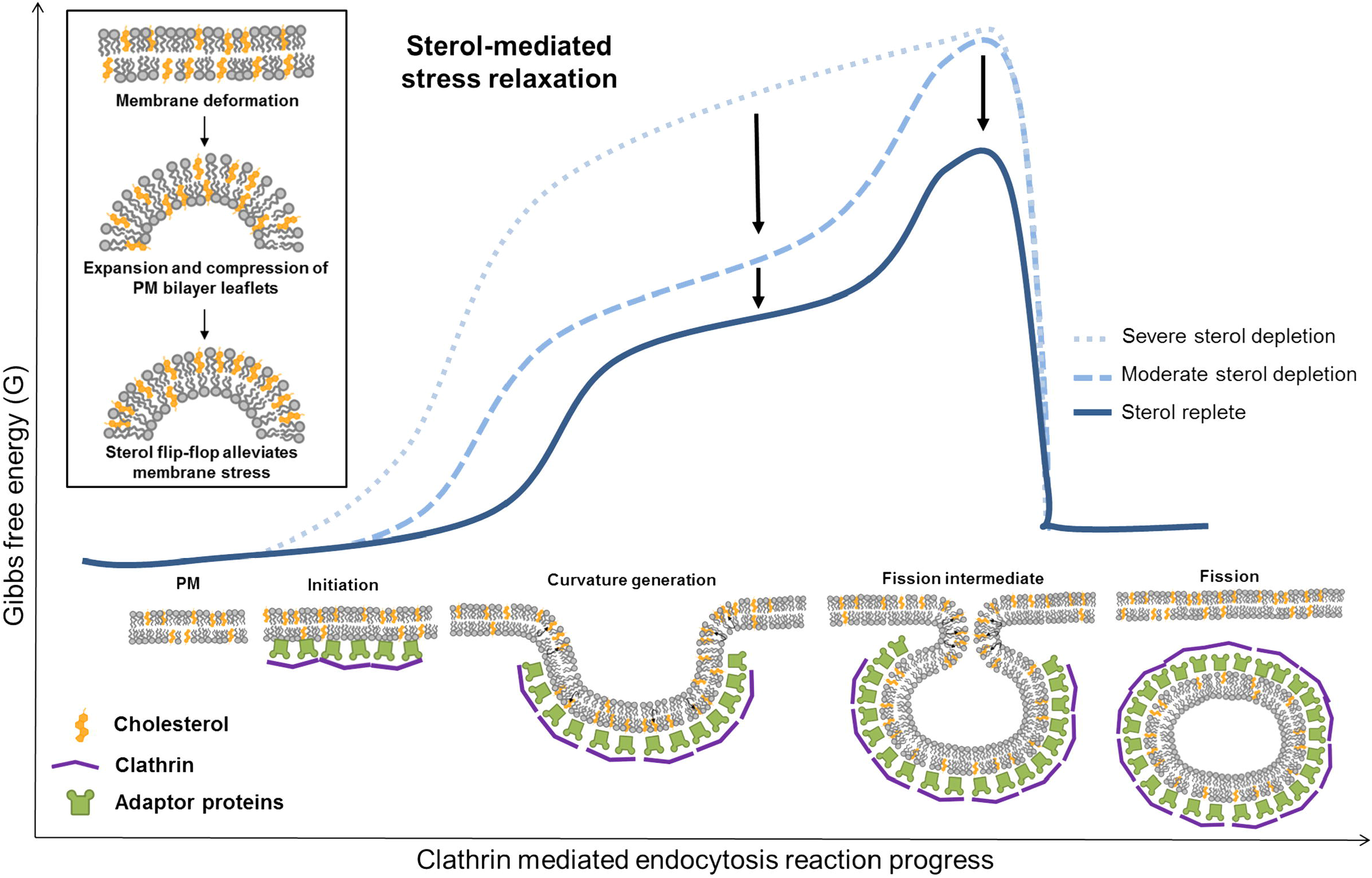
Model of cholesterol-mediated stress relaxation during clathrin-mediated endocytosis. Proposed mechanism of sterol dynamics during membrane remodeling and associated energy landscape of local curvature generation during CME. As sterols influence the tight packing of lipid bilayers, rapid cholesterol interleaflet flip-flop may relieve compression and expansion of the PM leaflets to negate energy costs associated with deformation of the membrane [**inset**]. This cholesterol-mediated stress relaxation reduces the energetic requirements necessary for membrane remodeling. Experimental evidence presented herein supports the model that cholesterol-mediated stress relaxation may facilitate initiation of curvature generation as well as formation of the highly curved vesicle neck. Tight lipid packing of sterols, thought to underlie formation of *L_o_* and *L_d_* domains (e.g. lipid rafts), may serve a driving force for membrane budding as increasing curvature minimizes line tension between phase boundaries.

In addition to modifying membrane bilayer architecture directly, sterol homeostasis may also impact CME through secondary mechanisms. Membrane curvature itself may influence the intrinsic activity of enzymes involved in membrane traffic (Bozelli et al., 2018). Furthermore, recruitment of curvature-sensing structural proteins may be affected, including ENTH/ANTH and BAR domain proteins which further promote PM curvature through hydrophobic insertion into the membrane or scaffolding mechanisms (Haucke and Kozlov, 2018). Cholesterol’s influence on lateral packing and ordering of the local lipid environment may also disrupt PI(4,5)P_2_ organization (Kwik et al., 2003), which may affect association of PH domain containing CME proteins. Packing of cargo is likely less affected by sterol depletion, as previous studies have demonstrated that the concentration of transferrin receptor (TfR) increased in accordance with coated pit surface area (Rodal et al., 1999). Though not required for CME in mammalian cells, actin assembles during the final stages of CME where it is thought to aid in the transition of domed to spherical shaped pits and vesicle scission under conditions where membrane tension is increased (Boulant et al., 2011; Hassinger et al., 2017; Skruzny et al., 2012). While disruption of actin integrity did not appreciably change CME responses to disruption of sterol synthesis, acute cholesterol depletion with MβCD is known to lead to stabilization of the actin cytoskeleton and increase membrane-cytoskeleton adhesions through micropipette aspiration and optical tweezer membrane tether experiments (Khatibzadeh et al., 2012). Given that the balance between endocytosis/exocytosis and the cytoskeleton are highly coordinated through membrane tension (Diz-Munoz et al., 2013), a holistic understanding of membrane surface dynamics in response to cholesterol depletion may lead to a more cohesive understanding of cellular responses and those observed within SLOS (Jiang et al., 2010).

As our findings favor a biophysical rather than a biochemical explanation for cholesterol’s role in facilitating membrane curvature, we would predict the findings presented here for CME may extend to other endocytic and membrane remodeling processes. Indeed, several other endocytic pathways, including caveolar, clathrin-independent, macropinocytosis, and clathrin- and caveolae/caveolin1-independent endocytosis, are known to be sensitive to cholesterol depletion (Doherty and McMahon, 2009). Within our own TEM analysis, we observed flattened and disassembled caveolar structures, consistent with reports that caveolin-1 may both sense and induce membrane curvature by cholesterol clustering (Krishna and Sengupta, 2019). Cholesterol also appears to be necessary for fusion pore kinetics during exocytosis as observed in platelets (Ge et al., 2010), chromaffin cells (Wang et al., 2010), and neurons (Najafinobar et al., 2016). In the limited studies where endocytosis and exocytosis have been measured simultaneously, sterol depletion appears to principally impair endocytosis (Subtil et al., 1999; Yue and Xu, 2015). Furthermore, intracellular recycling of TfR to the cell surface also appeared to be unaffected, suggesting the PM may be more susceptible to sterol depletion than other intracellular compartments (Subtil et al., 1999). A plausible explanation for these phenomena may lie in the transbilayer asymmetry of PM sterol distribution and directionality of sterol flux as lipid gradients maintained across organelles have been proposed to provide a self-organizing directionality to both the secretory and endosomal pathways (Levental et al., 2020).

Our studies suggest that reduced endocytic activity may be an important contributor to the cellular phenotypes observed within SLOS, potentially providing a unifying explanation for some of the varied and highly tissue-specific abnormalities reported within SLOS. As endocytosis and exocytosis are functionally coupled, impairment of CME may be linked to trafficking abnormalities previously described within secretory granules of the pancreas, pituitary and adrenal glands in *Dhcr7^−/−^* and *Sc5d^−/−^* mouse models (Gondre-Lewis et al., 2006). Similarly, impaired endocytic traffic may lead to dysregulation of the endosomal-lysosomal pathway which may be related to both aberrant Sonic Hedgehog signaling (Blassberg and Jacob, 2017; Blassberg et al., 2016) and compromised phagosome maturation (Futter et al., 2012; Ramachandra Rao et al., 2018) described within SLOS models. During embryonic development, CME is necessary for maintenance of pluripotency through regulation of pluripotent and differentiation signals (Narayana et al., 2019). Impaired Wnt/β-catenin and cadherin-related signaling was recently identified within SLOS patient-derived stem cells which display precocious differentiation (Francis et al., 2016), suggesting a possible role for CME deficits within cell fate decisions and developmental pathologies. Within the nervous system, dynamic communication between neurons is highly dependent upon intact vesicular trafficking and CME for rapid neurotransmission, receptor desensitization, dendritic spine plasticity, and myelination (Murthy and Stevens, 1998; Saheki and De Camilli, 2012; Watanabe et al., 2014; Winterstein et al., 2008). Interestingly, impairment of CME through *DNM1* mutations causes epileptiform discharges (Dhindsa et al., 2015), a prevalent finding in SLOS individuals (Schreiber et al., 2014). Rapid endocytosis is also of significant relevance to the brush boarder of small intestinal enterocytes whose microvilli exhibit high curvature and are highly enriched in cholesterol (Hansen et al., 2003; Huttner and Zimmerberg, 2001). As such, endocytic deficits could contribute to feeding difficulties and gastrointestinal intolerance commonly observed in SLOS. Previous studies have also observed increased levels of Tfn protein present within the retina of AY9944-treated rats (Tu et al., 2013) and cerebrospinal fluid isolated from SLOS subjects (Cologna et al., 2016), possibly due to reduced TfR-mediated internalization. The varied and complex role of cholesterol in different cellular functions indicates that malformations and functional deficits observed within disorders of cholesterol synthesis could be due to competing and disparate effects of the lipid. As such, the extent to which impairment of CME contributes to disease pathogenesis and clinical phenotypes observed will require additional careful study.

Current treatment strategies for SLOS include empirical dietary cholesterol supplementation, which has been shown to normalize plasma cholesterol with variable outcomes on sterol precursors (Linck et al., 2000). However, cholesterol supplementation does not universally correct plasma cholesterol concentrations nor does plasma cholesterol necessarily reflect tissue sterol content. Furthermore, the benefits of dietary cholesterol supplementation have been modest(Svoboda et al., 2012) and longitudinal studies indicate that baseline cholesterol levels are a better predictor of developmental trajectory than age of supplementation or increase in cholesterol serum levels (Sikora et al., 2004). Depressed CME activity may be linked to suboptimal efficacy of cholesterol supplementation by further reducing cholesterol bioavailability. Our studies indicate that once CME is effectively inhibited by AY9944, recovery of clathrin trafficking requires upwards of 6-12 hours to resume in the presence of FBS, yet recovers within minutes if cholesterol is delivered directly to the PM via MβCD carrier. Interesting, a secondary defect in LDL metabolism has been described within SLOS fibroblasts, where decreased LDL apolipoprotein degradation products were observed (Wassif et al., 2002). While this mechanism likely involves impaired NPC1 function(Wassif et al., 2002), it’s possible that a defect in LDL internalization due to impaired CME efficiency also contributes to this effect. A potential strategy to overcome impaired internalization and utilization of cholesterol *in vivo* may be direct delivery of cholesterol via a mechanism independent of receptor-mediated endocytosis, such as a cyclodextrin (CD) carrier. This seemingly straightforward approach could preferentially target PM properties likely mediating CME impairment as well as bypass impaired lysosomal function, necessary for liberation of cholesterol esters. Furthermore, our sterol substitution studies indicate that cholesterol precursors are more readily mobilized by MβCD, with cholesterol preferentially retained. Thus, cyclodextrin-mediated cholesterol delivery may have the added benefit of aiding clearance of sterol precursors. CDs have been widely studied for various applications within pharmacology (Vecsernyes et al., 2014). Most recently “empty” 2-Hydroxypropyl-β-Cyclodextrin (HPβCD) received regulatory approval for treatment of Niemann-Pick Type C1, currently undergoing Phase II and III clinical trials (Ory et al., 2017). A reversal of this strategy as a cholesterol delivery mechanism in SLOS could theoretically overcome several challenges hindering current cholesterol supplementation therapy.

In summary, we find that maintenance of cholesterol homeostasis is essential for efficient clathrin mediated endocytosis, demonstrate an active role for sterols in mediating membrane remodeling during this process, and provide mechanistic detail into the sterol functional requirements necessary for this process. While the extent to which altered CME dynamics and function contribute to the pathogenesis of disorders of cholesterol metabolism is unclear, further investigation into CME’s role within these orphan disease is warranted.

## ACKNOWLEDGEMENTS

This study was supported by NIH grants (NIGMS P20 GM103620 and P20 GM103548). A.D.H was supported by the National Science Foundation (#0953561), a National Science Foundation/EPSCoR Cooperative Agreement (#IIA-1355423), and the State of South Dakota through BioSNTR, a South Dakota Research Innovation Center. K.A.S and J.W.T. was supported by the Intramural Research Program of the National Heart Lung and Blood Institute, National Institutes of Health (ZIA HL006098). R.H.A was supported by the National Institutes of Health under a Ruth L. Kirschstein Fellowship (F30 NS106788). M.M.S. was supported by the Sanford Program for Undergraduate Research (P20 GM103620). We would like to thank the University of South Dakota Center for Brain and Behavior Research for supporting this project. We also thank Kelly Graber with the Sanford Imaging core and Maudi Killian with the Sanford Flow Cytometry core for technical expertise and use of imaging and sorting instruments. We also thank the NHLBI electron microscopy facility. Any opinions, findings, and conclusions or recommendations expressed in this material are those of the author(s) and do not necessarily reflect the views of the National Science Foundation or the National Institutes of Health.

## AUTHOR CONTRIBUTIONS

Conceptualization, R.H.A. and K.R.F.; Methodology, R.H.A., K.R.F., A.D.H., B.L.S., J.G.K.; Investigation, R.H.A., M.M.S., K.A.S., B.L.S., and E.M.B.; Software, H.V. and B.L.S; Writing— Original draft, R.H.A; Writing—Review & Editing, R.H.A, K.R.F., A.D.H, B.L.S, E.M.B, K.A.S., J.W.T; Funding Acquisition, R.H.A. and K.R.F.; Supervision, K.R.F., A.D.H, and J.W.T.

## MATERIALS AND METHODS

### Cell culture

HEK293T (parental and hCLTA^EN^-Tq2) and human SK-MEL-2 (hCLTA^EN^-Tq2 hDNM2^EN^-eGFP, as previously described(Scott et al., 2018)) were maintained in DMEM (Gibco, 4.5 g/L glucose, 110 mg/L pyruvate), supplemented with 10% (v/v) FBS (Hyclone), and 1,000 U/mL penicillin/streptomycin. For inhibition of cholesterol synthesis, cells were rinsed twice in serum-free DMEM and cultured under cholesterol deplete conditions in 7.5% delipidated serum (LPDS) for 48 h with small molecule inhibitors AY9944 (2.5 μM, Cayman Chemical; *DHCR7* inhibitor), U18666A (20 nM, Cayman Chemical; *DHCR24* inhibitor), Atorvastatin (1 μM, Cayman Chemical), or Simvastatin (Cayman Chemical), or at concentrations as otherwise specified. Acute sterol depletion was achieved by 1 h incubation with 5 mM MβCD (Alfa Aesar, 1303.31 g/mol) at 37°C. De-identified fibroblasts cultured from skin punch biopsies from Smith-Lemli-Opitz subjects (kind gift from Dr. Forbes Porter, NICHD) were maintained in DMEM (Gibco, 4.5 g/L glucose, 110 mg/L pyruvate), supplemented with 15% (v/v) FBS (Hyclone), and 1,000 U/mL penicillin/streptomycin. SLOS fibroblasts were rinsed twice in serum-free DMEM and cultured for 10 days in 7.5% (v/v) LPDS to induce biochemical profiles.

### CRISPR/Cas9 gene editing of clathrin light chain

C’ terminal targeting of the *CLTA* gene to enable fluorescence tracking with mTuroquise2 (mTq2) was performed using a homology-directed repair strategy as previously described (Anderson et al., 2018; Scott et al., 2018). Briefly, a guide RNA (5’-GCAGATGTAGTGTTTCCACA-3’) targeting the open reading frame in the immediate vicinity of the stop codon was cloned into the Cas9 expression vector pX330-U6-Chimeric_BB-CBh-hSpCas9 (gift from Feng Zhang (Cong et al., 2013), Addgene plasmid #42230). The donor vector consisted of 1 kB homology arms flanking a cassette containing puromycin N-acetyl-transferase expressed in-frame with *CLTA* and Tq2 via the self-cleavable peptide sequence P2A. 1 × 10^6^ HEK293T cells were transfected with 4 μg donor and 2.5 μg pX330 Cas9 plasmid (CalPhos™ Mammalian Transfection kit, Clontech). After 72 h, Tq2 positive cells were isolated with a FACSJazz Cell Sorter (BD Biosciences) and characterized as described previously (**Figure S1**) (Anderson et al., 2018).

### Preparation of delipidated serum

To facilitate cell culture under defined lipid conditions, fetal bovine serum (FBS) devoid of sterols, triglycerides, and other neutral lipids was prepared based on previously described techniques (Cham and Knowles, 1976). To prevent oxidation due to trace peroxides, 0.1 mg ethylenediamine tetraacetate (EDTA) was added per 50 mL FBS. Serum was mixed 1:2 with organic phase (consisting of 3:2 ratio diisopropyl ether:n-butanol) followed by vigorous stirring for 1 h protected from light. After centrifugation at 4°C for 15 min at 2200 rpm, the aqueous layer was collected and lyophilized. Following resuspension in molecular grade H_2_O, ITS-G supplement (Invitrogen) was added, the solution filter-sterilized, and stored at −20°C. GC/MS analysis indicated nearly undetectable cholesterol levels. Delipidated serum (LPDS) was substituted for regular FBS to stimulate expression of de novo cholesterol biosynthesis.

### Sterol loading of MβCD

Methyl-β-cyclodextrin (MβCD)-cholesterol complex commercially available (Sigma, C4951). For MβCD-sterol complexation, 50 mg/mL sterol stocks in 1:1 chloroform:methanol were prepared for desmosterol, lathosterol, cholestenone, cholesterol sulfate, 4β-hydroxycholesterol (Avanti Polar Lipids), 7DHC and cholesterol (Sigma or Avanti Polar Lipids). Stoichiometry of MβCD-cholesterol complexes typically consists of one cholesterol molecule entrapped within two CDs (1:2 molar ratio). We found optimal loading conditions using a 1:7 molar ratio, where desired sterol concentration was dried under nitrogen flow, followed by addition of 5 mM MβCD (Alfa Aesar, 1303.31 g/mol) prepared in serum-free DMEM. Sterol suspensions were sonicated for 5 min and incubated at 37°C with agitation overnight. Sterol crystals were subsequently removed via 0.45 μm filtration. Sterol loading rescue experiments were performed by incubation for 1 h at 37°C supplemented with 0.5% BSA, followed by removal and addition of 7.5% LPDS for imaging or subsequent analysis to limit prolonged exposure to MβCD.

### GC/MS sterol analysis

Cell pellets flash frozen on dry ice were reconstituted in 1 mL of water and lysed by successive freeze/thaw cycles in a 50°C bead bath. 50 μL per sample was removed for protein quantification (MicroBCA Protein Assay, Pierce Biotechnology). 1 mL saponification buffer containing 7% KOH in 92% ethanol with 10 ug/mL coprostan-3-ol (Abcam) was added to 900 μL cell lysate. Following saponification at 60°C for 1 h, an additional 1 mL of water was added to each sample and the aqueous phase extracted with 3 mL ethyl acetate by vortexing and centrifugation at 2200 rpm. The organic phase was then extracted with 2 mL water, concentrated to dryness by heating at 50°C under nitrogen flow, residue dissolved in 50 μL pyridine (CHROMASOLV Plus, Sigma), and sterols derivatized in 50 μL N,O-bis(trimethylsilyl) trifluoroacetamide with 1% trimethylchlorosilane (BSTFA + 1% TMCS, Thermo Fisher TS-38831) at 60°C for 1 h. Sterol levels were determined by GC/MS analysis as previously described (Kelley, 1995). Samples were analyzed by automatic injection of 1 μL of the derivatized sterol mixture into an Agilent 7890 GC using a split injection port (4 mm ID × 78.5 mm quartz wool liner, Restek 23309) leading to a 0.18 mm ID × 20 m 1,4-bis(dimethylsiloxy)phenylene dimethyl polysiloxane column (Restek, 43602). Helium was used as a carrier gas at a linear rate of 46.9 cm/sec. After 0.5 min at 170°C, the oven temperature was raised to 250°C at 18°C/min, then to 280°C at 3°C/min, and finally to 320°C at 20°C/min and held for 7 min. An Agilent 5977B mass spectrometer was operated in the electron impact mode at 70 eV with an ion source temperature of 275°C. Analysis was performed using MassHunter software. Identification of TMS ethers of natural sterols was determined through comparison to commercially available standards for cholesterol, 7DHC, lathosterol, and desmosterol (Avanti Polar Lipids, Inc.), as well as comparison to MS spectra through the National Institute of Standards and Technologies Standard Reference Database when available. Identification of 8DHC was inferred as an isomer of 7DHC and comparison to SLOS fibroblasts; zymostenol identification was based on spectra from Conradi-Hünermann-Happle syndrome (CDPXD2) fibroblasts. Retention times and mass to charge (*m/z*) ratios are summarized in **Table S1**. Representative MS fragmentation patterns available upon request. TMS derivatives of sterols exhibiting abundance <3% were excluded from analysis. For peak quantitation, sterol abundance was normalized to both the internal standard (coprostanol) and protein concentration. Data is presented relative to control samples using GraphPad Prism software.

### Transferrin uptake assay

HEK293T cells were plated onto 0.1% gelatin coated 12 mm coverslips or 96 well glass bottom plates (Cellvis) and treated for 48 h under respective treatment conditions. For AY9944 rescue experiments, cells were placed in DMEM/BSA (DMEM containing 0.5% w/v BSA; bovine serum albumin) alone or DMEM/BSA containing 5 mM MβCD-sterol complexes for 1 h in a 37°C humidified incubator containing 5% CO_2_. HEK293T cells were then incubated with transferrin (Tfn) conjugated to Alexa Fluor™ 555 (25 ug/mL, Invitrogen) for 30 mins at 37°C, followed by rinsing in ice-cold PBS, and fixation in 4% paraformaldehyde for 20 min. For high-content imaging analysis, 24 well glass bottom plates (Cellvis) were imaged using CX7 Rescue High-Content Screening (HCS) Navigator software (PerkinElmer, Waltham, MA) and standard HCS imaging protocol.

Patient fibroblasts were plated onto 0.1% gelatin coated 12 mm coverslips 3 days prior to imaging. Uptake was performed as described above using Tfn conjugated to Alexa Fluor™ 555 (25 ug/mL, Invitrogen) for 15 min at 37°C. Following incubation, cells were rinsed with ice-cold PBS. Cells were fixed in 4% paraformaldehyde for 20 min, rinsed three times in PBS, and mounted onto glass slides with Dako mounting media (Agilent). Cell outlines were traced using the freehand selection tool and the integrated density of 25 cells from 8-12 fields was quantified using ImageJ software (v1.8.0).

### Confocal microscopy

Images were captured using a Nikon A1R resonant scanning multispectral confocal microscope (Nikon Instruments, Inc. Melville, NY), equipped with a live cell chamber (37°C with humidified 5% CO_2_), Nikon Ti Perfect Focus system, and NIS-Elements analysis software (Nikon). 48 h prior to live-cell imaging, HEK293T cells plated onto 0.1% gelatin-coated FluoroDish™ culture plates (35 mm, World Precision Instruments Inc.) were rinsed twice with DMEM (Gibco) to remove residual lipids and respective treatments were added. Prior to imaging, culture media was refreshed. Time-lapse videos were obtained at 100x magnification (1.45 NA oil objective) at 5-20 s intervals over a 5 min duration.

### Polarized TIRF microscopy

SK-MEL-2 hCLTA^EN^-Tq2 hDNM2^EN^-eGFP cells were seeded onto fibronectin-coated (Corning) 25 mm coverslips at a density of 8.0 × 10^4^ cells/cm^2^ and imaged within 4-6 hours of plating. To facilitate membrane labeling, a fresh DiI (1,1’-Dioctadecyl-3,3,3’,3’-teramethylindocarboncyanine perchlorate) solution containing 1 μg/mL DiI in 2.5% DMSO/HBSS (Corning) was prepared for each coverslip from a 1 mg/mL DiI stock dissolved in DMSO and stored under nitrogen. After incubating the freshly prepared DiI solution at 37°C for 5 min, 200-300 μL was added dropwise to coverslips containing 1 mL HBSS. Cells were labeled for 20 s with gentle pipetting, rinsed three times in HBSS, and placed into imaging buffer. Imaging buffer for all treatment conditions consisted of Leibovitz’s L-15 media supplemented with either 10% FBS or delipidated (LPDS) serum (Hyclone) to monitor basal CME events. Following DiI labeling, coverslips were immediately imaged for no more than 45 min. To ensure consistency in DiI labeling, only cells with mean S polarization intensities between 350-800 were included in the analysis.

As previously detailed, TIRFM images were captured on a custom built Till iMic (Till Photonics, Germany) inverted microscope outfitted with a 60x/1.49 NA oil immersion objective and environmental chamber kept at 37°C (Scott et al., 2018). In brief, excitation for mTurquise2, eGFP, and polTIRF was achieved by 445 (470/22), 488 (510/10), 561 (595/50) nm lasers respectively, collected on three EMCCD cameras (iXon3 885, Andor Technology). Back focal plane centering was performed daily to optimize the incidence angle using MatLab analysis to map the angles of TIRF reflectance and determine mirror angle adjustments to center the optical plane. 2-point TIRF illumination facilitated polarization of the 561 nm laser to enable monitoring of membrane curvature. Focused excitation at 90° and 270° positions preferentially excite vertically oriented, DiI-labeled membrane (P polarization), while excitation at positions 0° and 180° selectively excite horizontally labeled membrane (S polarization).

### Polarized TIRFM image analysis

Image registration was performed to align Turquoise2 and eGFP channels onto the polTIRF channel. Calibration images of fluorescent beads on a glass coverslip were acquired simultaneously on all three detectors and used to determine the geometric transformation during pre-processing. Bias images, acquired by capturing images with a closed shutter, were subtracted from each frame of the raw data. Using MatLab, detection and tracking of fluorescence over time was performed using a modified version of cmeAnalysis software (Aguet et al., 2013) and custom scripts were used for association and classification of mTq2, eGFP, and P/S events (Scott et al., 2018). Detection criteria included a clathrin peak SNR > 4, dynamin SNR >2, clathrin lifetime >7 seconds, and a buffer of two consecutive frames before and after each clathrin track. Faulty tracks were filtered from the analysis due to clathrin events crossing paths with another track and curvature signal unrelated to the tracked clathrin event itself.

### Transmission electron microscopy

SK-MEL-2 cells were prepared as described for polTIRF imaging by seeding onto fibronectin-coated 25 mm coverslips and allowed to adhere for 5 h. Coverslips were transferred into 2 mL stabilization buffer (70 mM KCl, 5 mM MgCl_2_, 3 mM EGTA, 30 mM HEPES, pH 7.4) prior to unroofing with a 10 mL syringe and 22 gauge, 1.5 in needle containing 2 mL 4% PFA (Electron Microscopy Sciences) diluted in stabilization buffer. Coverslips were transferred to fresh 2% PFA/stabilization buffer and allowed to fix at RT for 20 min. Coverslips were mounted in 2% glutaraldehyde and sealed with VALAP (1:1:1 vasoline, lanolin, paraffin) for shipping. Sample preparation and TEM imaging of platinum cell membrane replicas was performed as previously described (Sochacki et al., 2017; Sochacki et al., 2014). Representative full EM images available upon request.

### Statistical analysis

All statistical analysis was performed using GraphPad Prism 8.0.2 (GraphPad Software, Inc., CA, US). Homogeneity of variances was tested by the Brown-Forsythe test. In cases of equal variances, data was analyzed by one-way ANOVA and post hoc Dunnett t-test for multiple comparisons relative to LPDS control group. In cases of unequal variances, Welch ANOVA and post-hoc Dunnett’s T3 test was performed. P◻<◻0.05 was accepted as significant. Statistical details for each experiment can be found within figure legends.

### Data availability

Datasets generated in this study are available from the corresponding authors upon request.

### Code availability

Programming generated in this study is available from the corresponding authors upon request.

## SUPPLEMENTAL INFORMATION TITLES AND LENGENDS

**Video 1. Confocal video of HEK293T hCLTA-mTq2^EN^ cells demonstrating immobilization of clathrin trafficking at the PM shown in Figure 1**. Endogenous clathrin trafficking in HEK293T cells cultured in 7.5% LPDS without **[Panel 1]** or with addition of AY9944 **[Panel 2].** Rescue of clathrin trafficking following sterol depletion with AY9944 by addition of cholesterol-loaded MβCD (MβCD-Chol) **[Panel 3]**. Confocal images taken at midplane. Acquisition: ~8s/frame. Frame rate: 15 frames/s.

**Video 2. Confocal video of HEK293T hCLTA-mTq2^EN^ cells demonstrating rescue of clathrin trafficking deficits by delivery of cholesterol or 7DHC via MβCD carrier as shown in Supplemental Figure 2.** Comparison of endogenous clathrin trafficking in HEK293T cells cultured in 7.5% LPDS **[Panel 1]** or following sterol depletion with 48 h AY9944 treatment **[Panel 2]**. AY9944 induced clathrin trafficking deficits could be rescued by sterol reconstitution with cholesterol-loaded MβCD (MβCD-Chol) **[Panel 3]** or 7DHC-loaded MβCD (MβCD-7DHC) **[Panel 4]**. Confocal images taken at midplane. Acquisition: ~16s/frame. Frame rate: 15 frames/s.

**Video 3. polTIRFM time-lapse video of SK-MEL-2 hCLTA-mTq2^EN^ hDNM2-eGFP^EN^ cell shown in Figure 3**. Clathrin, dynamin, and membrane curvature dynamics within 10% FBS control cell. hCLTA-Tq2, Dynamin2-eGFP, and P/S ratio indicating positive curvature generation pseudocolored cyan, green, and magenta respectively. Acquisition: 2s/frame. Frame rate: 20 frames/s.

**Video 4. TIRFM time-lapse videos of SK-MEL-2 hCLTA-mTq2^EN^ hDNM2-eGFP^EN^ cells under altered sterol conditions as shown in Figure 3**. hCLTA-Tq2 pseudocolored cyan, Dynamin2-eGFP pseudocolored green. Acquisition: 2s/frame. Frame rate: 20 frames/s.

