## Supplementary Materials for "Sterols lower energetic barriers of membrane bending and fission necessary for efficient clathrin mediated endocytosis"

| Sterol, sterol derivatives | Retention Time (min) | Molecular Ion *m/z* (Da) | Base peak *m/z* (Da) | Spectral Ions *m/z* (Da) |
|---|---|---|---|---|
| Cholesterol sulfate | 10.1 – 10.6 | Not detected | 368 | 353, 260, 247, 147 |
| Coprostanol TMS | 12.5 - 12.6 | Not detected | 370 | 403, 355, 215 |
| Cholesterol TMS | 13.7 - 13.8 | 458 | 329 | 443, 368, 353, 129 |
| 8-dehydrocholesterol TMS | 13.9 - 14.0 | 456 | 351 | 441, 366, 325 |
| Zymostenol TMS | 14.1 - 14.2 | 458 | 458 | 443,353,355,229,213 |
| Desmosterol TMS | 14.2 - 14.3 | 456 | 129 | 441, 366, 343, 327 |
| 7-dehydrocholesterol TMS | 14.3 - 14.6 | 456 | 351 | 441, 366, 325 |
| Lathosterol TMS | 14.6 - 14.7 | 458 | 255 | 443, 353, 229, 213 |
| Cholestenone | 15.1 - 15.2 | 384 | 124 | 369, 299, 229 |
| 4β-hydroxycholesterol TMS | 15.5 – 15.6 | Not detected | 366 | 456, 441, 417, 327 |

**Supplemental Table 1. Chromatographic and mass spectral parameters of sterols measured by GC/MS analysis.** Sterols isolated from cultured cells were derivatized to their respective trimethylsilyl (TMS) ethers to facilitate separation by gas chromatography on a 20 m, 0.18 mm internal diameter Rxi-5 Sil column (Restek, 43602). Retention times indicated above. Coprostanol was used as an internal standard in all experiments for normalization. Major fragment ions (*m/z*) are noted above. Sterols isolated from cultured cells were derivatized to their respective trimethylsilyl (TMS) ethers to facilitate separation by gas chromatography on a 20 m, 0.18 mm internal diameter Rxi-5 Sil column (Restek, 43602). Retention times indicated above. Coprostanol was used as an internal standard in all experiments for normalization. Major fragment ions (*m/z*) are noted above. Related to **Figures 1, 2, and 5**.

| **Sterol** | **Hydrophobicity*** (XLogP3) | **Polarity**** TPSA (Å²) | **Ordered lipid domain (raft) stability** | **References** |
|---|---|---|---|---|
| Cholesterol | 8.7 | 20.2 | ++ | Tempo quenching in small unillamellar vesicles (SUVs) [1]. 1:1:1 Cholesterol:Sphingomyelin:DOPC containing giant unilamellar vesicles (GUVs) induce positive curvature at phase separation boundaries by confocal and fluorescence correlation spectroscopy (FCS) [2]. |
| Desmosterol | 8.3 | 20.2 | + | Tempo quenching in SUVs [1]. |
| 7-dehydrocholesterol | 8.0 | 20.2 | +++ | Tempo quenching in SUVs [1]. 7DHC containing liposomes and SLOS rat model brains contain detergent-resistant domains [3]. |
| Lathosterol | 8.3 | 20.2 | +++ | Tempo quenching in SUVs [1]. |
| Cholestenone | 8.4 | 17.1 | - | Cholestenone does not support phase separation in 1:1 DOPC: sphingomyelin GUVs by confocal fluorescence microscopy at any concentration [4]. |
| 4ß-hydroxycholesterol | 7.4 | 40.5 | (-) | Polar hydroxyl group positioned within the hydrophobic region of the bilayer predicted to destabilize packing of lipid domains [2]. |
| Cholesterol sulfate | 8.2 | 72.0 | - | Pure cholesterol sulfate containing GUVs do not induce phase separation; Quaternary mixtures of DOPC/SM/cholesterol sulfate/cholesterol GUVs exhibit small, bell-shaped $L_o$ domains with negative curvature at phase boundaries [4]. |

*XLogP3: Computed octanol/water partition coefficient.
**Topological Polar Surface Area (TPSA): Computed surface sum over all polar atoms in a molecule.

**Supplemental Table 2. Physical properties and phase separation behavior of sterols used to assess the steroidal structural requirements for CME.** Structurally disparate sterols were chosen based on differences in ring and hydrocarbon tail saturation, predicted interleaflet translocation (flip-flop) dynamics, and the ability to support phase separation in model membranes. Ordered lipid ($L_o$) domain/raft stabilizing (+) or disrupting (-) sterols indicated relative to cholesterol (++). Predicted () and supporting experimental evidence are summarized under ‘references’. Hydrophobicity and polarity measures published on PubChem. DOPC, 18:1 (Δ9-Cis) phosphocholine; 7DHC, 7-dehydrocholesterol; SLOS, Smith-Lemli-Opitz syndrome. Related to **Figure 5**.

| Subject | Cell line | Mutations | Clinical Phenotype | Biopsy age | Gender |
|---|---|---|---|---|---|
| n/a | BJ | None | Unaffected | newborn | M |
| SLOS-066 | MN | p.M1V/p.Q98X | Mild SLOS | 2 years | M |
| SLOS-029 | CWI | p.T93M/c.964-1G>C | Classical SLOS | 6 months | F |
| SLOS-098 | A2 | p.964-1G>C homozygous | Severe SLOS | 1 day | M |

**Supplemental Table 3.** General characteristics of control and Smith-Lemli-Opitz syndrome patient fibroblasts utilized in this study. Cell lines were described previously [5, 6].

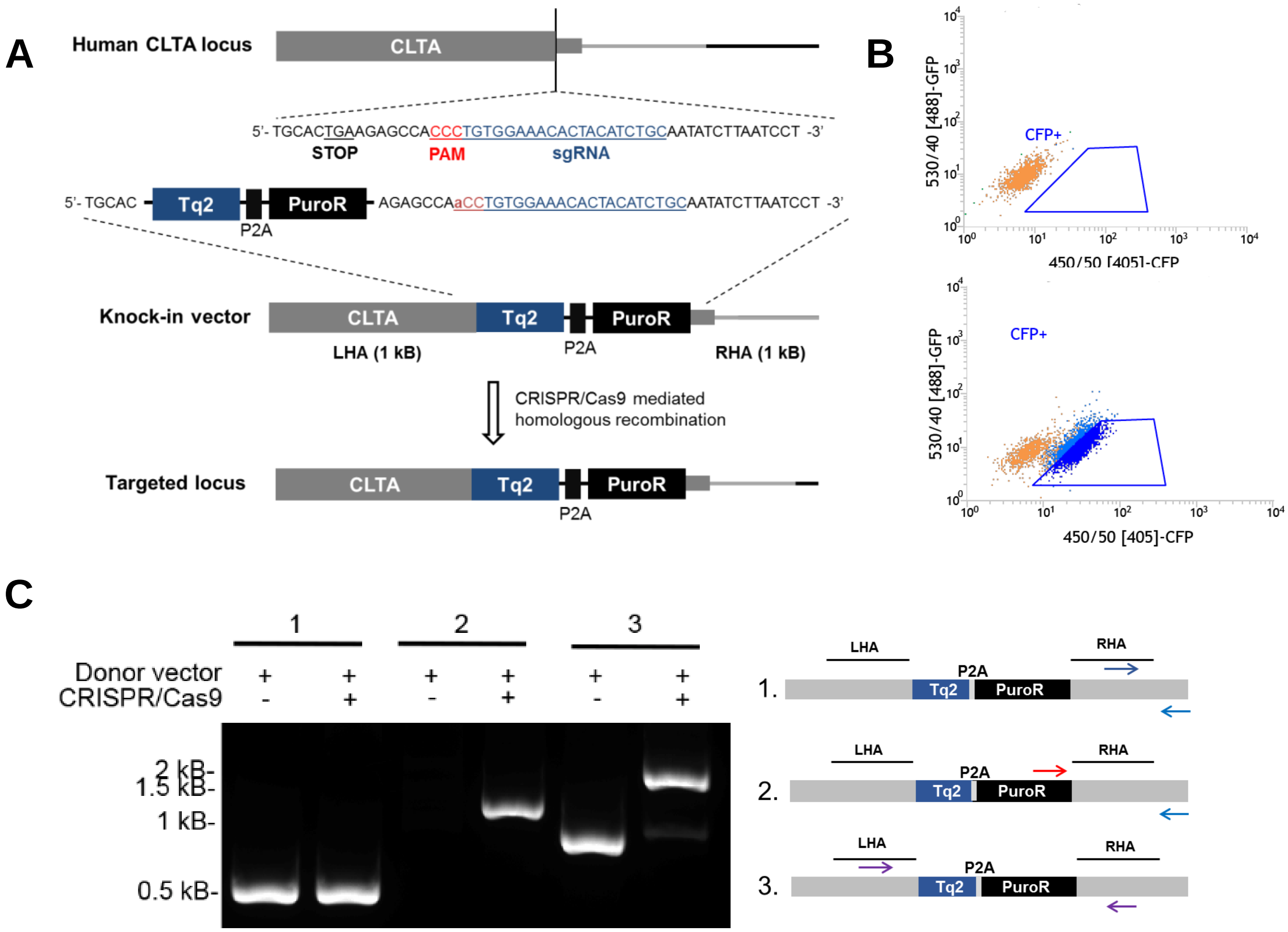


**Supplemental Figure 1. Characterization of HEK293T hCLTA-Tq2$^{EN}$ cell line**. **A)** Schematic of CRISPR/Cas9 mediated knock-in strategy targeting the C-terminus of clathrin light chain A (*CLTA*) locus in frame with mTurquoise2 (Tq2), followed by a self-cleavage peptide sequence (P2A) and puromycin resistance (PuroR). Homology arms flanking the Turquoise2-P2A-PuroR cassette were amplified from genomic DNA and inserted into a pDONOR2 vector by Golden gate assembly simultaneously using BsaI sites. To prevent Cas9 recognition after integration, the PAM recognition sequence within the donor vector was mutated. **B)** Selection of edited HEK293T cells by flow cytometry. Sorting of HEK293T transfected with only the donor vector to control for non-selective integration **[Top]** compared to donor and pX330 gRNA targeting the *CLTA* locus **[Bottom]. C)** PCR demonstrating targeted insertion of mTq2 into the CLTA locus utilizing primer pairs 5'- TTGCTTGCCAGTGTCCCTCAGTTTA -3' / 5'-GCTTTAATAATTGCTTGGAACATCACCT-3' (template control, lanes 1-2); 5'-TACGAGCGGCTCGGCTTCA -3' / 5'- GCTTTAATAATTGCTTGGAACATCACCT-3' (Homology-directed repair, lanes 3-4); 5'- ATCTTGGGAAAGCCAGAATGTCATT -3'/5'-CCTAAACTGAGGGACACTGGCAA-3' (allelic insertion, lanes 5-6). Related to **Figures 1, 2**.

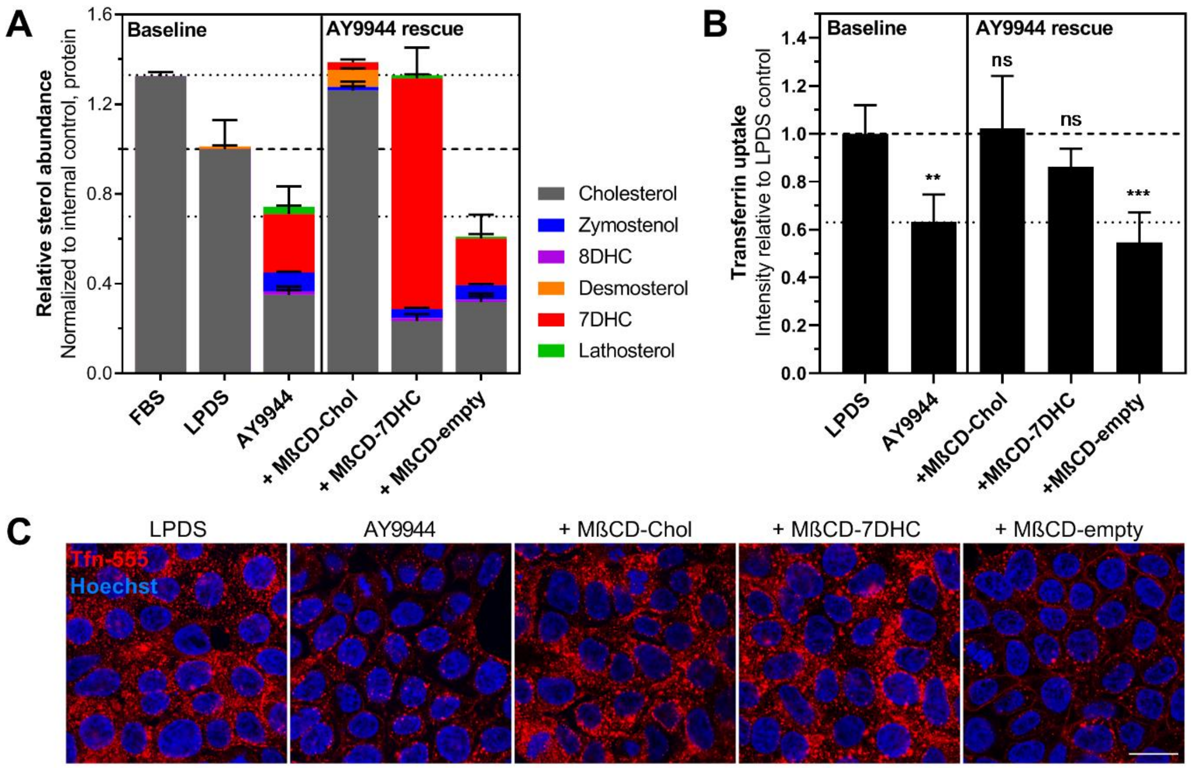


**Supplemental Figure 2. 7-dehydrocholesterol can compensate for cholesterol in facilitating clathrin mediated endocytosis.** Inhibition of CME due to sterol depletion by AY9944 treatment can be robustly rescued by direct delivery of cholesterol or 7-dehydrocholesterol (7DHC) via MβCD carrier. **A)** Sterol profiles of AY9944 treated HEK293T cells following 1h incubation with sterol loaded MβCD (Mean ± SD). N = 2 biological replicates from independent experiments. **B)** Tfn uptake relative to controls cultured in 7.5% LPDS for 48 h (Mean ± SD). **, $P < 0.01$; ***, $P < 0.001$; one-way ANOVA ($F(4,20) = 11.89$ , $p < 0.0001$) and Dunnett's test versus LPDS control (N = 5 biological replicates from 3 independent experiments, ~1,500 cells per replicate). **C)** Representative confocal images taken mid-plane following 30 min incubation with AF-555 conjugated transferrin (Tfn). Scale bar = 20 µm.

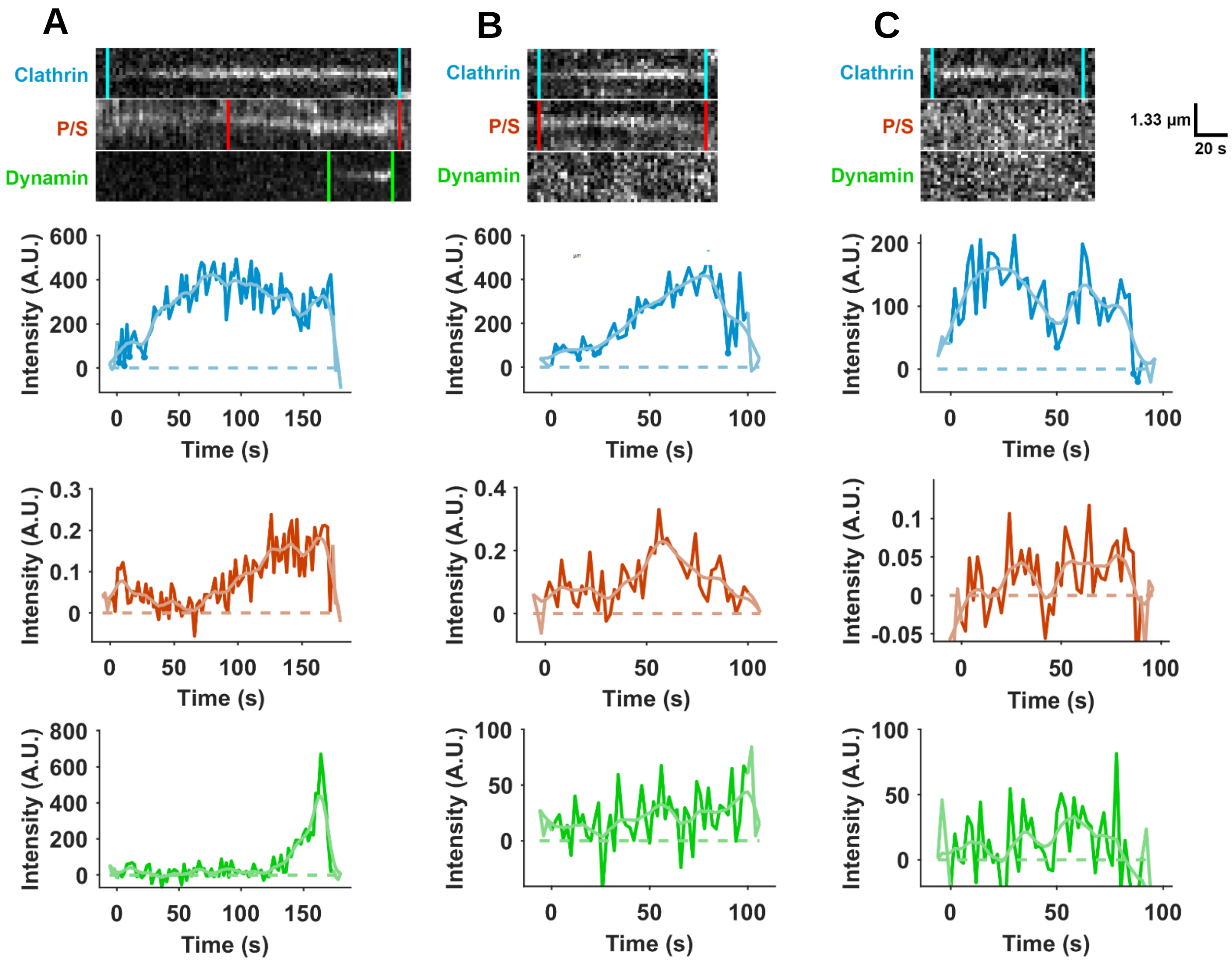


**Supplemental Figure 3. Additional polTIRF intensity tracings of clathrin events.** Kymographs generated from the center pixel of each tracked event with the lifetime of each component indicated by vertical bars. Background corrected intensity tracings for each component are below. Clathrin-Tq2 signal is shown in cyan, P/S curvature signal in red, and dynamin-GFP in green. Individually tracked clathrin events: **A)** valid CME event, **B)** clathrin event with positive curvature generation but lacking dynamin recruitment, and **C)** flat clathrin event with neither P/S or dynamin recruitment. Related to **Figure 3H**.

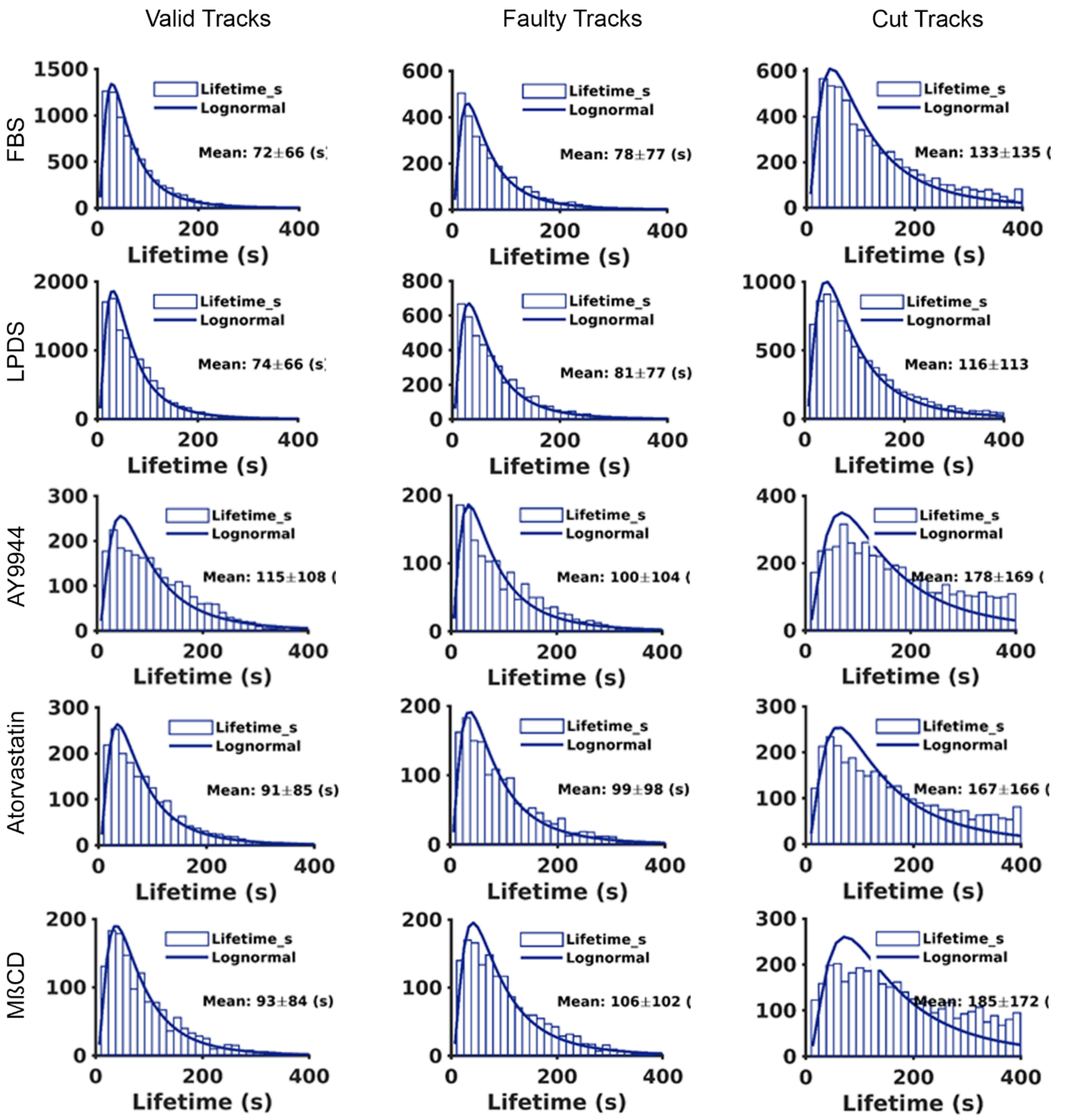


**Supplemental Figure 4. Clathrin lifetime distribution by track categorization**. Frequency of clathrin lifetimes observed in SK-MEL-2 (hCLTA-Tq2$^{EN}$/hDNM2-eGFP$^{EN}$) cells. Valid tracks are considered for CME productivity. Known lifetime of cut tracks extend before or beyond the first and last frames. Faulty tracks were excluded from further analysis. Related to **Figure 3F, 3G**.

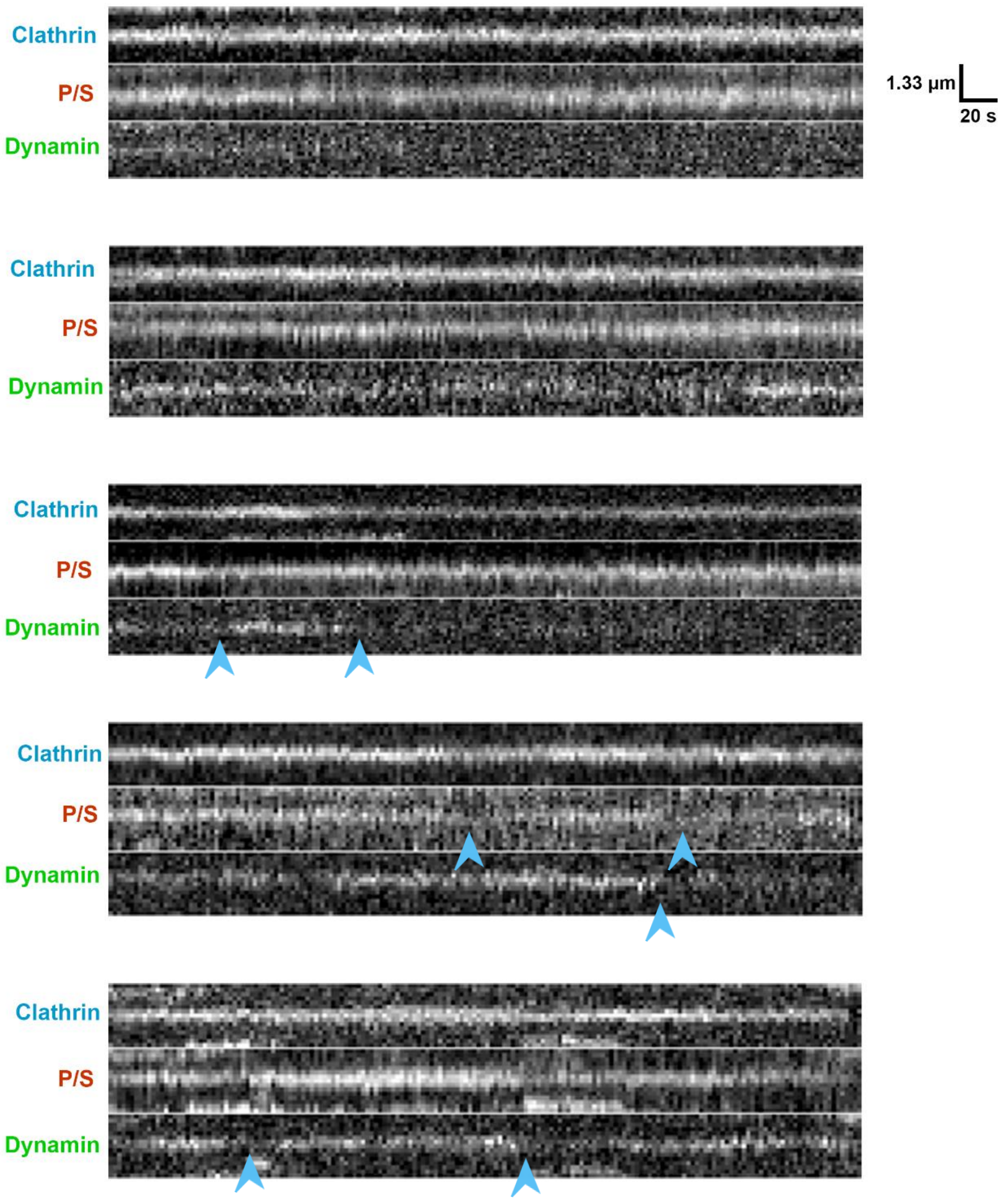


**Supplemental Figure 5. Representative kymographs of persistent clathrin tracks.** Kymographs generated from the center pixel of single tracked events observed in SK-MEL-2 (hCLTA-Tq2$^{EN}$/hDNM2-eGFP$^{EN}$) cells following sterol depletion with 2.5 µM AY9944 for 48 h. Note variation in curvature signal and dynamin association along static clathrin tracks (arrowheads). Related to **Figure 3**.

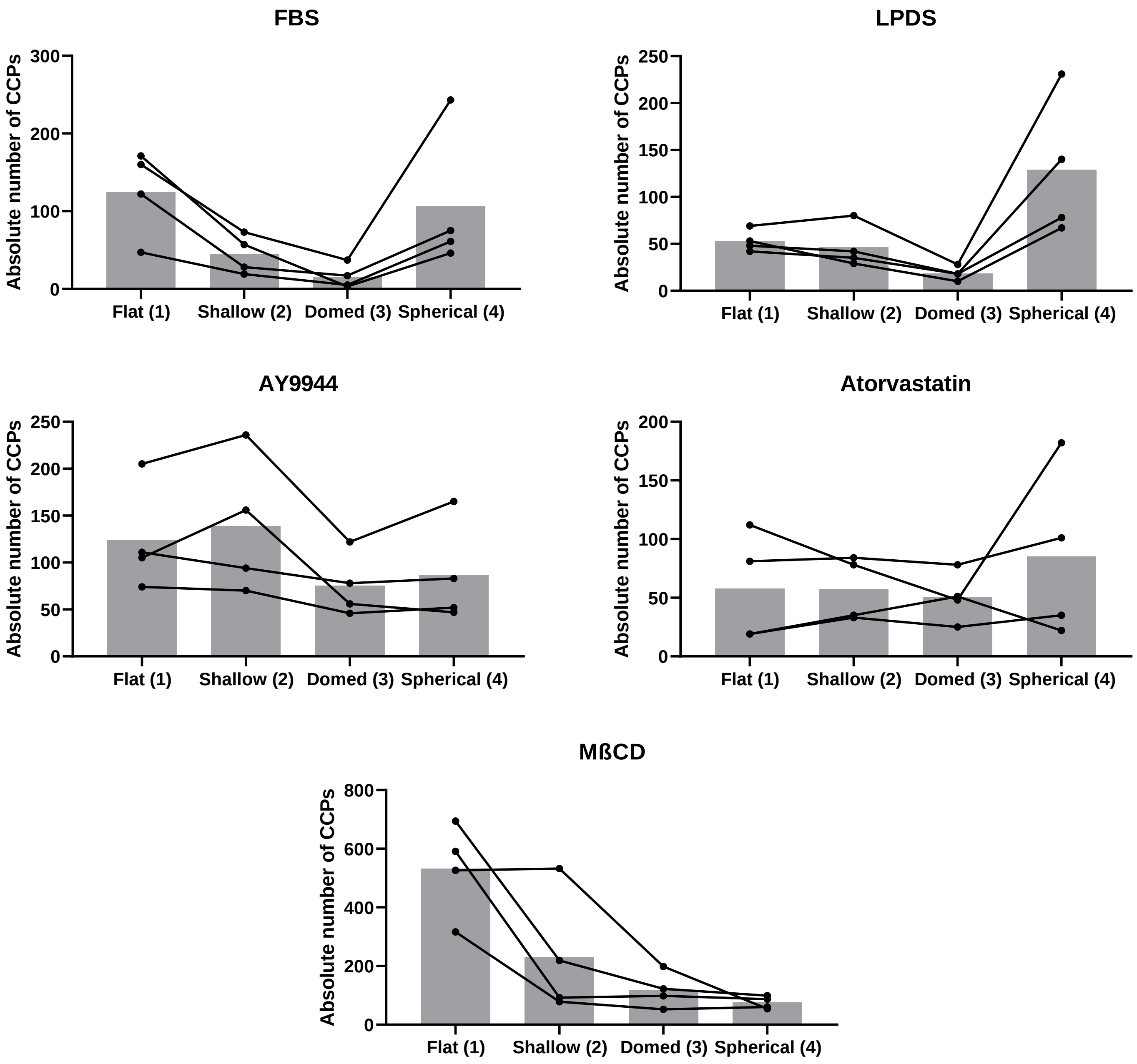


**Supplemental Figure 6. Distribution of curvature classification amongst TEM replica analyses.** Absolute number of clathrin structures analyzed per unroofed SK-MEL-2 cells by curvature classification. Distribution of clathrin structures within one replica is indicated by each line (total of 4 cell replicas per condition from 2 independent experiments were analyzed). Representative wide-view of platinum replicas available upon request. Related to **Figure 4A, 4B**.

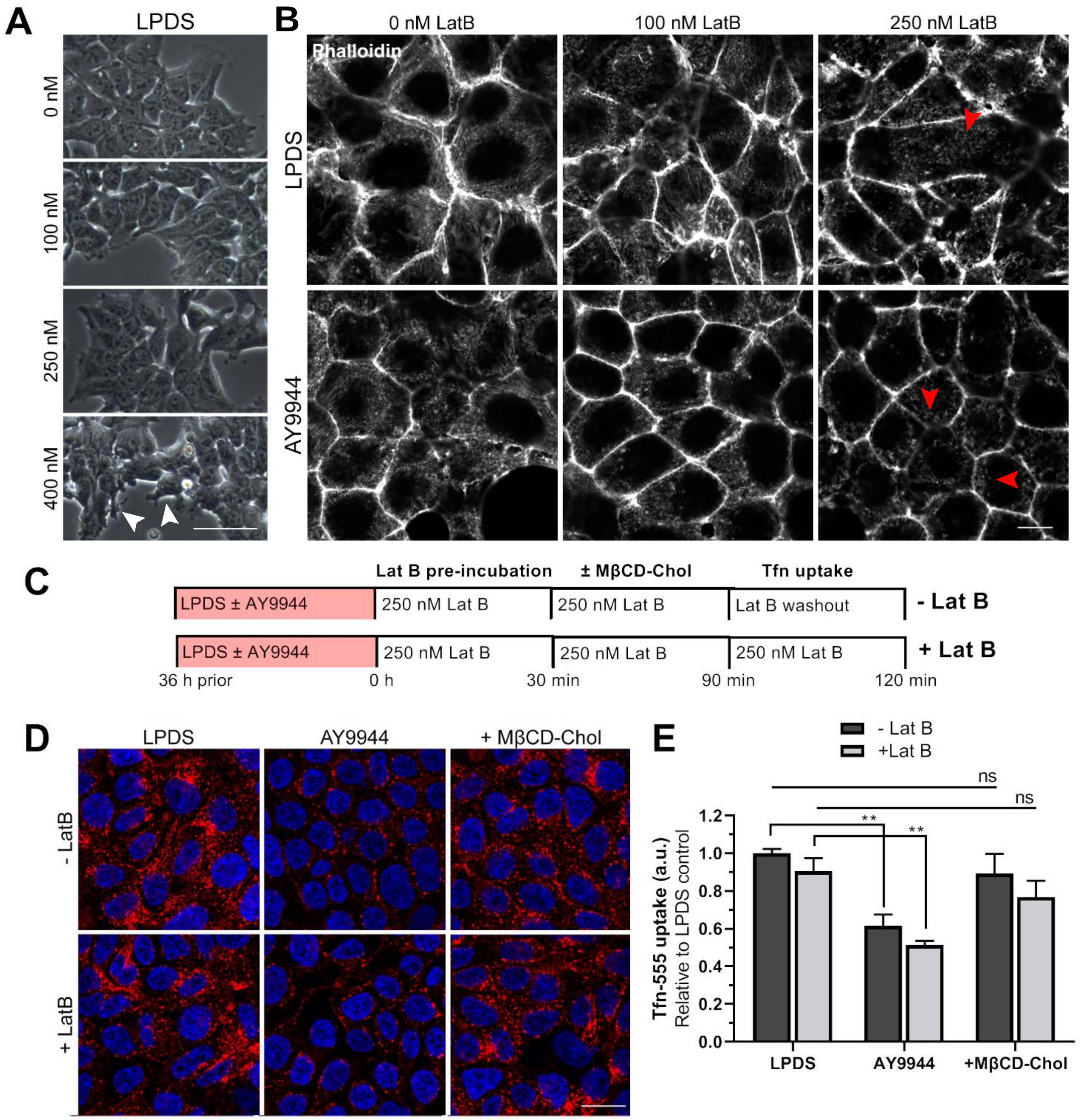


**Supplemental Figure 7. Actin network integrity not required for rescue of CME deficits secondary to sterol depletion.** Validation of Latrunculin B (Lat B) treatment in HEK293T cells optimized for a 2 h incubation period to access actin dependence of cholesterol-dependent CME activity. **A)** Low concentrations of Lat B (50-250 nM) did not cause retraction or gross cell morphology changes following a 2 h treatment by time-lapse DIC microscopy, as observed at higher concentrations (white arrowheads). Scale bar = 50 µm. **B)** Actin organization in LPDS and LPDS + AY9944 (DHCR7 inhibitor) treated cells following Lat B treatment for 2 h. Treatment with 250 nM Lat B disrupted most of the delicate apical actin network, leaving bright actin punta (red arrowheads). Scale bar = 10 µm. **C)** Schematic of experimental design for Lab B treatment and washout in relation to rescue of sterol content by addition of MβCD-Cholesterol prior to transferrin (Tfn) uptake assay to access CME function. **D)** Representative confocal images taken mid-plane following 30 min incubation with AF-555 conjugated Tfn. Scale bar = 20 µm. **E)** Compared to washout controls (- Lat B), Lat B treatment trended toward diminished Tfn uptake in all groups; however, Lat B pre-treatment did not prevent rescue of Tfn internalization following direct correction of cholesterol cellular content by MβCD sterol delivery, suggesting that cholesterol's influence over CME activity is independent of actin dynamics. Tfn internalization relative to controls grown in 7.5% LPDS for 36 h (Mean ± SD). **, $P < 0.01$; ns, not significant; one-way ANOVA ($F(5, 18) = 7.360$, $p = 0.0006$) and Tukey's test versus LPDS controls (N = 4 biological replicates from 2 independent experiments, ~1,500 cells per replicate).
